# Visuomotor association orthogonalizes visual cortical population codes

**DOI:** 10.1101/2021.05.23.445338

**Authors:** Samuel W. Failor, Matteo Carandini, Kenneth D. Harris

## Abstract

Training in behavioral tasks can alter visual cortical stimulus coding, but the precise form of this plasticity is unclear. We measured orientation tuning in 4,000-neuron populations of mouse V1 before and after training on a visuomotor task. Changes to single-cell tuning curves were apparently complex, including appearance of asymmetric and multi-peaked tuning curves. Nevertheless, these changes reflected a simple mathematical transformation of population activity, suppressing responses to motor-associated stimuli specifically in cells responding at intermediate levels. The strength of the transformation varied across trials, suggesting a dynamic circuit mechanism rather than static synaptic plasticity. This transformation resulted in sparsening and orthogonalization of population codes for motor-associated stimuli. Training did not improve the performance of an optimal stimulus decoder, which was already perfect even for naïve codes, but the resulting orthogonalization improved the performance of a suboptimal decoder model with inductive bias as might be found in downstream readout circuits.

---

Visual stimuli trigger patterns of activity across a multitude of neurons in the visual cortex. These patterns of activity can change following task training, and these changes persist even when the stimuli are presented outside the task context ^1–17^. Studies of representational plasticity have mainly analyzed changes in the tuning of single cells, but the nature of these changes appears complex and potentially contradictory. For example, some studies find increases^1,8,11^ and others decreases^4,12,16^ in the numbers of neurons representing task stimuli; some find broadening^6^ and others sharpening^3,5,9,15^ of orientation tuning curves; others find diverse types of tuning curve change such as asymmetric shifts or slope increases ^13,17^. This apparent complexity might arise at least in part from differences in the exact methods to select cells for analysis and to quantify their selectivity.

An alternative approach is to consider representational plasticity at the population level. Indeed, the population response to a stimulus defines a representation in a high-dimensional vector space, similar to the high-dimensional representations constructed by machine learning algorithms^18–20^. If it were possible to mathematically summarize the effects of task training on these population responses, this might help harmonize the diverse reported effects of training on single-cell tuning, providing clues to their biological mechanism and function.

A common hypothesis for the representational plasticity observed with task training is that it increases the fidelity of stimulus coding. Cortical responses vary between repeated presentations of an identical stimulus, and this variability could limit the ability of even an ideal observer to decode the stimulus from neuronal activity^21–23^. It has been suggested that task training changes the size and correlation structure of trial-to-trial variability, thereby improving the fidelity of the population code over that found in naïve cortex^1,7–9,11,12,24^. This hypothesis, of course, presupposes that the population code in naïve cortex does suffer from low fidelity. Although this may be true for some stimuli, the representations of oriented gratings – a stimulus often used in learning experiments – are extremely reliable^25^. Additionally, in some studies of somatosensory cortex and olfactory bulb, training did not improve the fidelity of neuronal responses^26,27^. Thus, if population responses to visual stimuli do change with learning, these changes may serve a function other than noise reduction.

A second hypothesis for the function of representational plasticity in sensory cortex pertains not to the abstract optimal decoder, but rather to the kind of suboptimal decoder that might be found in reality, e.g., in downstream brain structures. Even without noise, learning systems exhibit “inductive biases”, meaning that they learn some types of stimulus-response associations more readily than others, which may in turn be shaped by “priors” on the type of associations likely to be encountered^28–31^. Animals are likely to produce similar behavioral reactions to sensory stimuli evoking similar neural representations, and to more readily distinguish stimuli evoking different representations^28,29,32,33^. For an animal to learn different associations to two stimuli, the cortical representations of the stimuli should become differentiated^34^, with the firing vectors they evoke becoming more orthogonal. These changes would make the inductive bias in the downstream brain structures more suited to the task, allowing them to more readily associate the different representations with different actions. Amongst the diverse types of single-cell plasticity reported in visual cortex, two effects –reduction in the number of responsive cells and tuning-curve sharpening – are consistent with a reduction in overlap between populations responding to different stimuli^17^, and thus with population-level orthogonalization of stimulus representations.

We used two-photon calcium imaging to study how the tuning of V1 populations changes after mice learn to associate opposing actions with two oriented gratings. Stimulus coding was already perfectly reliable in naïve animals, so could not be improved further by task training. When analyzed at a single-cell level, plasticity appeared highly complex, with individual neurons’ tuning curves transformed in multifarious ways including previously reported sharpening and asymmetric slope change, as well as novel phenomena such as development of multimodal tuning curves. Nevertheless when viewed at a population level, these diverse single-cell phenomena could be quantitatively explained by a simple nonlinear transformation of population rate vectors, by a function whose convexity is largest for motor-associated stimuli. The strength of transformation varied consistently across the population on a trial-by-trial basis, suggesting that it emerges from circuit dynamics rather than static changes in the circuit as might be expected from synaptic plasticity. This transformation caused the population responses to become more orthogonal, an effect that was strongest for the stimuli with opposite motor associations, and improved the performance of a model decoder exhibiting inductive biases such as might be found in downstream brain circuits.

## Results

We trained mice in a visuomotor association task (Fig. 1A-B; Supplementary Fig. 1). Mice were shown pairs of grating stimuli and were trained to form motor associations with gratings of two orientations (45° and 90°) representing opposite motor contingencies (turn the wheel towards vs away), while a third orientation was a distractor (68°) that was presented with the same frequency as the motor-associated stimuli. No other orientations were presented during task performance.

**Figure 1.**
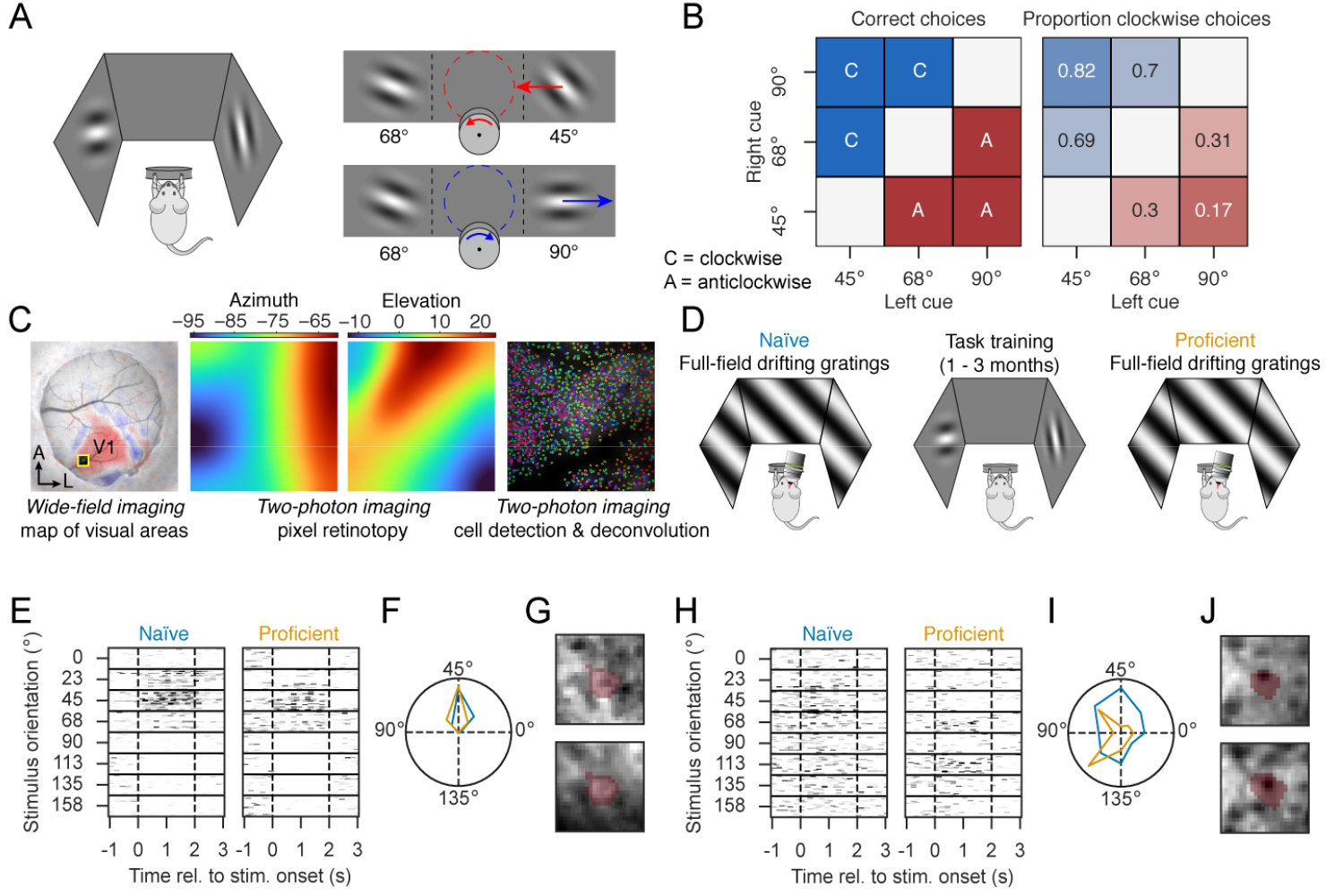
A visuomotor association task and two-photon calcium imaging methods. **A**, On each trial mice are presented with two stimuli and then turn a wheel to move them on the screens. Turning towards the 45° stimulus or turning away from the 90° stimulus yields a reward, but 68° stimuli are distractors. **B**, Correct choices for all stimulus pairings (left) and the average proportion of clockwise choices across mice taken from their ten highest performing sessions (right). **C**, Pipeline for imaging neural activity. Left: V1 was located using widefield imaging with sparse noise stimuli (red/blue: sign map; yellow outlined square: re-gion selected for two-photon imaging). Middle: retinotopy map for the two-photon field of view. Right: colored outlines of detected cells. **D**, Timeline of experiments. Responsesto drifting grating stimuli were recorded in naïve mice, and in the same mice after they had become proficient at the task. **E**, Raster representation of responses to repeated grating stimuli for an example cell in a naïve mouse, and the same cell when the mouse was profi-cient at the task. **F**, Orientation tuning curves of the same cell before and after training, superimposed in polar coordinates (radius represents mean response of the cell to each orientation). **G**, Mean intensity images of the region of V1 containing the ROI of the recorded cell, top: naïve, bottom: proficient. The cell’s ROI is shown as a red overlay. **H-J**, Same as **E**-**G** for a cell with weaker orientation selectivity.

To study how task training affected cortical representations of visual stimuli, we assessed the orientation tuning of excitatory cells in V1 using two-photon calcium imaging (Fig. 1C-D). We obtained two recordings: one before task training began (naïve condition) and one after training was complete (proficient condition). We recorded 5,041 ± 2,347 and 4,197 ± 1,486 cells in naïve and proficient mice (mean ± SD, n = 5 mice), with 277 ± 232 cells tracked between the naïve and proficient cases using a custom semi-supervised ROI matching algorithm (mean ± SD, n = 4 mice; Methods). In both training conditions, drifting gratings were presented in a passive session where no rewards were given, and the wheel was not coupled to visual stimuli. In these passive sessions we did not see wheel or body movements from either the wheel rotary encoder or the videographic data. Presentation of gratings caused pupil constriction, which was more prominent following training but not specific to any orientation (Supplementary Fig. 2A-B). It evoked minimal whisking that was not significantly affected by training or orientation (Supplementary Fig. 2C-D). Thus, even though body movements modulate visual cortical activity^35–38^, analyzing passive stimulus responses avoided this potential confound.

### An ideal decoder distinguishes task stimuli perfectly even before training

Individual cells showed a range of tuning characteristics and formed a population code that had extremely high fidelity in both naïve and proficient mice. Some neurons in both naïve and proficient mice showed sharp orientation tuning (Fig. 1E-G). Other neurons showed broader tuning, with multi-peaked tuning curves particularly noticeable in proficient mice (Fig. 1H-J). Applying dimensionality reduction to the population activity (Methods), we observed that population responses to different grating stimuli showed essentially no overlap (Supplemental Fig. 3A). As a first test of the fidelity with which V1 encoded grating orientation, we asked how reliably population activity distinguished the two motor-associated orientations (45° vs 90°) by training an optimal decoder (linear discriminant analysis; Fig. 2A). This yielded essentially 100% cross-validated accuracy for all orientations in both naïve and proficient mice (Fig. 2B; Supplementary Fig. 3B).

**Figure 2.**
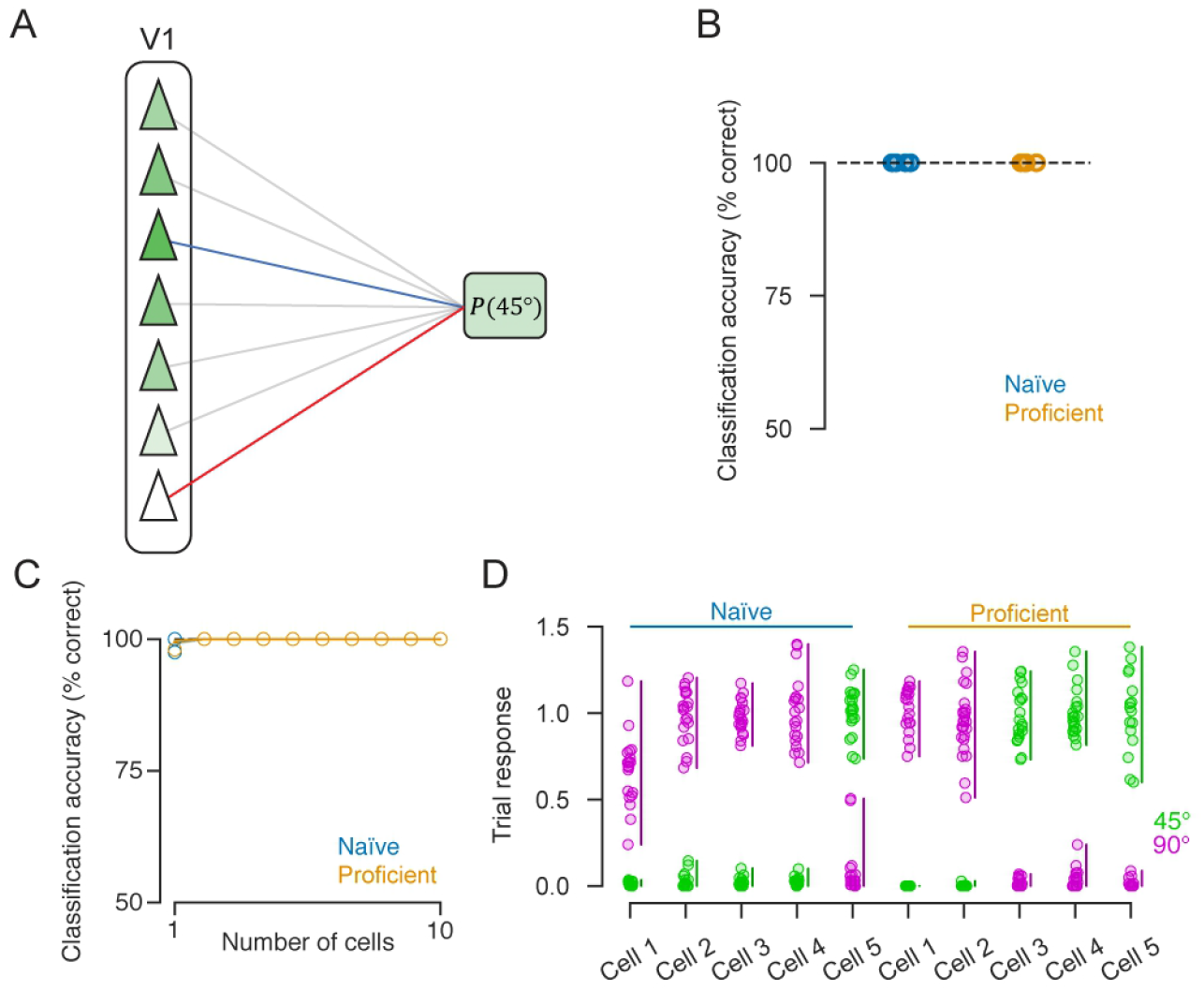
An ideal decoder performs perfectly even before training. **A**, Schematic of linear decoder. The decoder was trained to distinguish between 45° or 90° stimuli by discriminant analysis, which takes a weighted sum of measured cortical activity and applies a soft threshold to obtain a posterior probability for the stimulus orientation. **B**, Cross-validated classification accuracy for decoding stimulus orientation from naïve and proficient mice with linear discriminant analysis. Dashed line indicates perfect performance (n = 5 mice); shading for standard error is present but too small to see. **C**, Accuracy of decoding from an optimal subset of neurons, selected greedily from the population, as a function of number of subset size. No significant difference between naïve and proficient conditions was found. **D**, Single-trial responses of example cells, showing how the stimulus can bede-coded from just one cell’s activity. Each column shows one cell’s activity on all trials (left 5 columns, naïve conditions, right 5 proficient). Magenta circles show the cell’s activity on individual trials with 90° stimuli; green circles: activity on individual trials with 45° stimuli. Bars to the right of each column show ranges of response to the two stimuli, which are completely non-overlapping.

**Figure 3.**
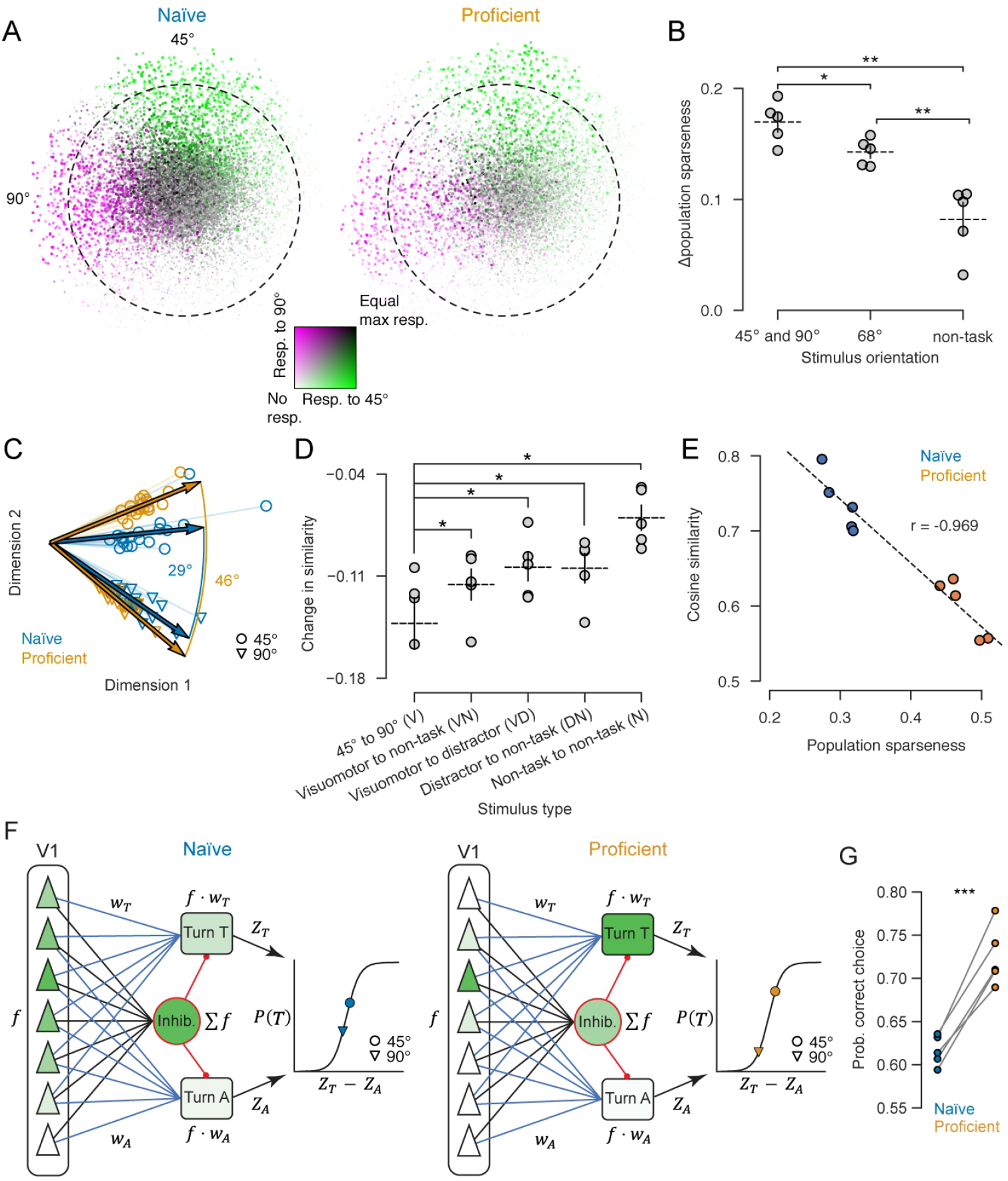
Task training sparsens and orthogonalizes cortical population codes. **A**, “Bull-seye plots” showing mean population responses to the motor-associated orientations 45° and 90°, for naïve and proficient conditions. Each point represents a cell (16,000 randomly-selected cells per plot), at a polar location determined by the cell’s mean orientation preference (angle) and orientation selectivity (distance). Color represents the cell’s response to the 45° (green) and 90° (magenta) stimulus orientations on an additive scale so points responding to both stimuli appear gray; the point’s size and brightness (light to dark) represents the cell’s maximal response to these two stimuli. Dashed circle is the threshold for considering a neuron highly selective (distance of 0.64). **B**, Change in population sparseness between naïve and proficient conditions, for each stimulus type. Error bars: mean and SEM (N = 5 mice). Significance stars: paired samples *t*-test. **C**, PCA projection of 45° (circle) and 90° (triangle) trials from one mouse in naïve (blue) and proficient (orange) condition. Arrows: normalized mean response vectors for each stimulus and condition. The angle between mean population responses to the two stimuli goes from 29° (naïve) to 46° (proficient). **D**, Change following training in cosine similarity between population responses to the two motor-associated stimuli, and two non-task stimuli. Error bars: mean and SEM (N = 5 mice); see Supplementary Fig. 7 for all stimulus pairs. Significance star: paired samples *t*-test. **E**, Population sparseness plotted against cosine similarity for naïve and proficient experiments (N = 5 mice, naïve and proficient conditions). **F**, Schematic for downstream decoder model with inductive bias. Cortical cells (triangles), whose activity is clamped to experimental measurements, project to two decision units (one for clockwise and one for anticlockwise turns) via excitatory weights (*w*_*e*_ and *w*_*A*_) proportional to the mean response of each presynaptic cell to the corresponding stimulus. The decision units also receive divisive feedforward inhibition, proportional to the unweighted sum of presynaptic excitatory activity. The difference between the firing rates of the two decision units is passed into a sig-moid function to obtain the probability of a turn toward vs away from the stimulus. Green shading: cartoon illustration of activity levels when cortical activity is dense in naïve mice (left), or sparse in proficient mice (right). Circle and triangle on logistic curve: model outputs when driven by recorded cortical responses to motor-associated stimuli. **G**, Model performance as assessed by probability of correct choice, for models driven by activity recorded of naïve and proficient mice (n = 5 mice). Significance stars: paired samples *t*-test.*, p < 0.05; **, p < 0.01.

This result does not support the hypothesis that correlated neural noise presents a fundamental limit to the fidelity of stimulus coding in naïve animals, at least for the stimuli used here. Because this hypothesis has been influential, we analyzed our contradictory evidence in detail (Appendix 1). These analyses revealed that training-related changes in stimulus representations do not effect the fidelity of stimulus encoding because a small population of neurons encoded the stimuli with extremely high reliability, in both naïve and proficient conditions. Indeed, training a sparse decoder by stepwise regression showed it was possible to decode the stimulus with 100% accuracy from the activity of just one of these neurons, in both naïve and proficient conditions (Fig. 2C,D). We did not observe a difference in the performance of an ideal decoder following training, even after handicapping the decoder in various ways, such as by limiting the number of neurons it could access (Appendix 1).

If downstream regions applied to V1 output the same optimal decoders as in these analyses, there would be no need for V1 to adapt its code to facilitate a visuomotor association. We therefore turned to our second hypothesis: that the changes in visual representation of the motor-associated stimuli would improve performance in a potentially suboptimal downstream decoder.

### Training sparsens and orthogonalizes population responses to motor-associated orientations

Consistent with the inductive bias hypothesis, we found that training differentiates population responses to the motor-associated stimuli by sparsening them and making them more orthogonal. To visualize changes in the population code, we developed a “bullseye plot” (Fig. 3A), to compare the responses of many neurons to two stimuli. We selected 16,000 cells (from 5 mice) randomly to equalize numbers between conditions, and for each plotted a point in polar coordinates with angle determined by the cell’s preferred orientation and radius by its orientation selectivity (distance). Point color represents the cell’s response to the two task stimuli using a two-dimensional colormap, with cells responding exclusively to 45° shown in green, cells responding exclusively to 90° in magenta, and cells responding to both in gray/black. Although the plots for naïve and proficient mice display the same number of neurons, fewer points are visible for the proficient mice (right), indicating that fewer neurons responded to the task stimuli (nonresponsive neurons are represented in white and thus invisible). This suggests that the population code grew sparser after task training. Moreover, fewer points were gray/black, indicating that fewer cells responded to both stimuli (visible as a reduction in the number of gray/black points), and thus an orthogonalization of the codes for the two stimuli. This effect was predominately due to neurons with low selectivity (gray/black neurons in the center) becoming unresponsive; highly selective neurons were generally unaffected.

To quantify changes in population sparseness, we used the Treves-Rolls population sparseness measure^39,40^, which quantifies the fraction of cells with near-zero activity to each stimulus (Methods; this differs from “lifetime sparseness”, which quantifies the fraction of stimuli driving each cell and will be discussed below). Population sparseness increased for the motor-associated orientations 45° and 90°, significantly more than for the distractor stimulus 68°, and in turn more than for non-task stimuli (Fig. 3B; 45° and 90° vs 68°: p = 0.04; 45° and 90° vs non-task: p = 0.008. 68° vs non-task: p = 0.003; paired samples *t*-test, n = 5 mice).

Previous work has reported that a sparsening of population responses can arise simply from passive stimulus exposure^27,41^. The motor-associated and distractor stimuli appeared with equal probability during training, but random fluctuations led to different exact numbers of times each stimulus was presented to each mouse; furthermore, the total number of presentations differed between mice due to differences in training duration. We tested for an effect of the number of stimulus exposures on population sparsening using ANCOVA analysis, finding a significant effect of stimulus orientation, but not of cumulative exposures (Supplemental Fig. 3C). Thus, simple stimulus exposure does not explain the higher sparseness observed for the motor-associated stimuli.

The increase in population sparseness occurred together with an orthogonalization of the population code. To visualize orthogonalization, we mapped population response vectors for the motor-associated stimuli into two dimensions using principal component analysis (Fig. 3C). To quantify this orthogonalization we computed the cosine similarity between the mean response vectors to the motor-associated stimuli. After training, this similarity decreased significantly more than the similarity between other stimulus pairs (Fig. 3D; 45° to 90° vs: informative to non-task, p = 0.036; informative to distractor (68°), p = 0.0215; distractor to non-task, p = 0.0256; non-task to non-task, p = 0.0136; paired samples *t*-test, n = 5 mice; see Supplementary Fig. 3D for stimulus type definitions and all pairwise similarity comparisons). The mice showing the greatest sparsening also showed the most orthogonalization (Fig. 3E): cosine similarity was correlated with population sparseness (p < 0.0001, mixed effects model; n = 5 mice), but showed no further dependence on training condition once sparsening was accounted for (p = 0.12, mixed effects model, n = 5 mice), suggesting that sparsening and orthogonalization are two reflections of a single process. Indeed, a correlation between sparsening and orthogonalization is expected for mathematical reasons: sparsening increases the number of near-zero components in the population response vectors, and thus decreases the cosine similarity between stimulus pairs^42^ (Appendix 2.2).

### A computational model for the benefits of sparsening and orthogonalization

We next addressed the question of how population sparsening and orthogonalization we observed might benefit decision making. We defined a computational model, focused not on the performance of an ideal decoder – which would perform perfectly in both naïve and proficient conditions (Fig. 2B) – but on a neural decoder with inductive bias related to the similarity of cortical representations. We hypothesized that orthogonalization could differentiate responses to the two task stimuli, by reducing a prior bias to respond similarly to them.

The modelled decision circuit receives input from a population of V1 neurons whose activity is clamped to that measured in individual trials before or after training (Fig. 3F). The brain location of this decision circuit is not relevant but for concreteness one might imagine the striatum, whose circuitry our model is based on. The decision circuit contains three units: one “decision unit” for each of the two choices (turn towards or turn away), and one feedforward inhibitory unit providing divisive inhibition. We use these three single units for simplicity; the brain of course would use three populations for each. The decision units receive feedforward excitation from each V1 neuron, with weights proportional to the V1 neuron’s response to the appropriate stimulus, as could be learned by a simple Hebbian rule. The inhibitory unit receives feedforward excitation from all cortical neurons with constant equal weights, and its output acts divisively on both decision units^43^. Finally, the probability of a turn towards the task stimulus is modeled as a logistic function of the difference between the two decision units^44^.

The model’s performance in the task was significantly better when the V1 activity was clamped to activity recorded in proficient conditions than naïve conditions (Fig. 3G). To understand why, we looked at how the activity of different model components changed with changes in cortical population sparseness. As expected, sparser cortical activity resulted in lower total excitatory input to the decision units (Supplemental Fig. 3E). However, it did not increase the difference between the excitatory inputs received by the two decision units (Supplemental Fig. 3F). This is because sparsening of cortical activity primarily reduces the activity of weakly tuned neurons (Fig. 3A), which feed into both decision units and so do not affect the difference in their activity. Rather, sparser cortical activity boosted the activity of the decision units by reducing feedforward inhibition (Supplemental Fig. 3G). Indeed, the output of the decision network is determined by the difference between the decision units’ activity, which equals the difference between their excitatory inputs divided by the amount of feedforward inhibition. Training-induced sparsening decreased the latter while leaving the former essentially unchanged, leading to an increase in the network’s output and therefore in the probability it would make the correct choice (Supplemental Fig. 3H). Thus, sparsening and orthogonalization of cortical activity causes the output of this simple readout circuit to become differentiated, reducing its prior bias to produce the same response to both stimuli.

### Training suppresses responses to task stimuli in weakly-tuned cells

Sparsening and orthogonalization are consequences of a change in the sensory population code, but do not fully characterize this change. To understand the precise changes in code structure, we next examined the mean responses of individual neurons to gratings of all orientations, summarized by their orientation tuning curves (Fig. 4A). In naïve animals, tuning curves often had a standard single-peaked profile. In proficient animals, however, tuning curves were often irregular and multipeaked. Multimodal tuning can be detected by comparing a cell’s modal orientation preference (i.e., the orientation that drives it most strongly; circles in Fig. 4A) to its circular mean orientation. The circular mean vector is a sum of unit orientation vectors weighted by the cell’s response to that orientation of grating (arrows in Fig. 4A); this vector’s direction is the cell’s mean orientation preference, and its magnitude the cell’s orientation selectivity. The modal and mean orientations are equal for unimodal symmetrical tuning curves but can differ when tuning curves are multimodal (Supplementary Fig. 4A-B). Comparing the modal and mean orientations of recorded neurons indicated that cells with multimodal tuning curves were more common in proficient mice; these cells typically had weaker orientation tuning, and their mean (but not modal) orientation was close to the motor-associated orientations 45° and 90° (Supplementary Fig. 4C).

**Figure 4.**
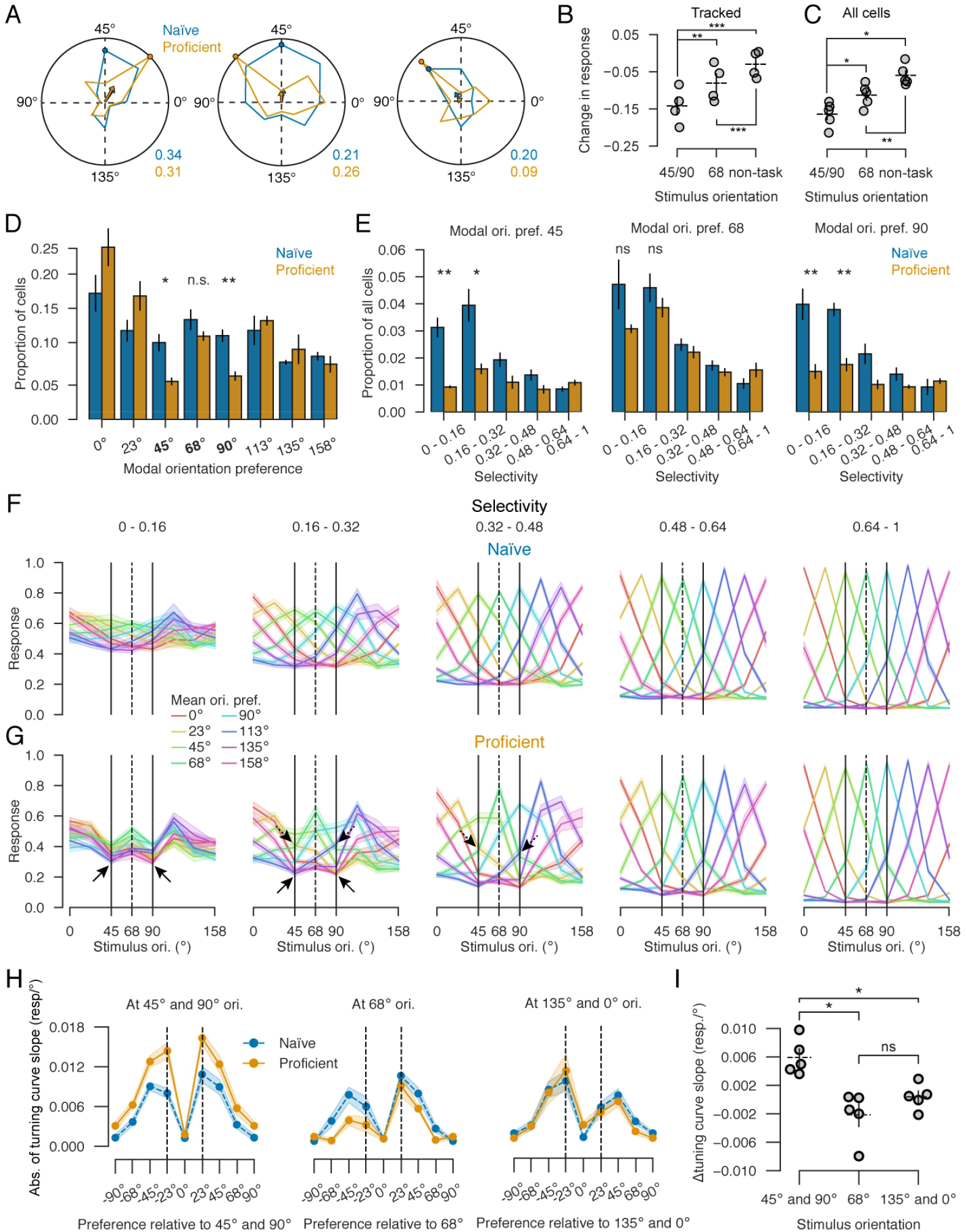
Training suppresses responses to task stimuli in weakly-tuned cells. **A**, Examples of naïve and proficient orientation tuning curves for three cells. Colored polar curves: mean response to each orientation; dots: response to modal orientation; arrows: circular mean vectors representing mean orientation preference (angle) and orientation selectivity (length). Numbers at lower right: naïve and proficient selectivity indexes. **B**, Average training-induced change in responses to the motor-associated, distractor, and non-task orientations for tracked neurons. Error bars: mean and SEM (N = 5 mice). Significance stars: hierarchical linear mixed effects model (n = 1107 cells, N = 5 mice). **C**, same for all recorded neurons. Error bars: mean and SEM (N = 5 mice). Significance stars: paired samples *t*-test (N = 5 mice). **D**, Proportion of cells with each modal orientation preference, in naïve and proficient mice. Error bars: SEM (N = 5 mice). Significance stars: paired samples *t*-test. **E**, Proportion of cell population that had modal orientation preference 45° (left), 68° (center), and 90° (right) and specified orientation selectivity. Significance stars: paired samples t-test. **F**, Average orientation tuning curves for cell groups defined by mean orientation preference (color) and selectivity (column) in naïve mice. Solid vertical lines indicate motor-associated orientations, dashed the distractor (68°). Y-scale measures mean deconvolved response of all cells in the group, scaled by maximal response (see Methods). Shading: SEM (N = 5 mice). **G**, Same plot for proficient mice. Solid arrows highlight suppression of cell responses to the motor-associated orientations 45° and 90°. Dashed arrows highlight tuning curve steepening. **H**, Tuning curve slope analysis, as in Ref. ^13^. The three plots show tuning curve slopes at motor-associated, distractor, and non-task orientations. Each point shows an average over cells grouped by mean preferred orientation relative to the orientation the slope is computed at (x-axis) and training condition (color). Shading: SEM (n = 5 mice). **I**, Change in tuning curve slope at motor-associated, distractor, and non-task orientations, averaged over cells with adjacent orientation preferences for each experiment (at dashed lines in **H**). Error bars: mean and SEM (N = 5 mice). Significance stars: paired samples *t*-test. *, p < 0.05, **, p < 0.01.

Closer examination of example cells (Fig. 4A) suggested that this bimodality occurred because responses to the motor-associated orientations 45° and 90° had been suppressed following training. This was confirmed by smaller responses to the motor-associated orientations 45° and 90° in proficient than naïve mice (Fig. 4B-C). The training-related suppression of responses to these task-associated stimuli was larger than to the distractor stimulus 68°, which was in turn more suppressed than responses to orientations that were not presented during training (Tracked cells, 45° and 90° vs 68°: p = 0.0007; 45° and 90° vs non-task: p < 0.0001; 68° vs non-task: p < 0.0001; hierarchical linear mixed effects model, n = 1107 cells, N = 5 mice. All cells, 45° and 90° vs 68°: p = 0.04; 45° and 90° vs non-task: p = 0.01; 68° vs non-task: p = 0.008. paired samples *t*-test, n = 5 mice).

The suppression of responses to task orientations led to a change in the distribution of modal orientation preferences, primarily in weakly-tuned cells (Fig. 4D-E). Task training significantly decreased the fraction of cells modally preferring the motor-associated orientations (45° and 90°), but not the distractor orientation (68°) (Fig. 4D; 45°: p = 0.014; 68°: p = 0.228; 90°: p = 0.006, paired-sample *t*-test, n = 5 mice). The decrease in cells modally preferring the motor-associated orientations came specifically from cells of low orientation selectivity (assessed by the length of the circular mean response vector; arrows in Fig. 4A): there was no decrease in the number of cells strongly tuned for motor-associated orientations (Fig. 4E; 45°: p = 0.005 and 0.037 for orientation selectivity 0 - 0.2 and 0.2 - 0.4; 68°: p = 0.130 and 0.390; 90°: p = 0.001 and 0.013, paired samples *t*-test, n = 5 mice). Analysis of cells tracked between naïve and proficient recordings confirmed that cells with weak preference for the motor-associated orientations in naïve mice were most likely to change their tuning preference following task training (Supplementary Fig. 5A-B).

Training also changed tuning curve shapes, in a manner dependent on a cell’s preferred orientation and selectivity (Fig. 4F-G). We divided the recorded cells into groups according to their orientation selectivity and mean orientation preference (mean preference was used rather than modal as it is more stable; Supplementary Fig. 4) and plotted the mean tuning curves of cells in each group before and after training, using held-out repeats. In naïve mice, tuning curves were unimodal and symmetrical, with similar shapes for all mean orientations (Fig. 4F). For proficient mice, however, a different structure appeared (Fig. 4G). Weakly tuned neurons were suppressed by the motor-associated orientations, even including cells whose mean orientation was motor-associated. Restricting the analysis to neurons tracked between naïve and proficient conditions gave similar results, confirming that responses to the motor-associated stimuli 45° and 90° were suppressed more strongly than both the distractor stimulus 68° (which was shown equally often during training) or stimuli that had not been shown at all (Supplementary Fig 5B-C). Suppression of motor associated orientations also led to an asymmetrical increase in tuning curve slopes at these orientations, specifically for neurons whose mean orientation preference flanked one of the task stimuli (Fig. 4H-I), as reported in primates^13^.

### A simple quantitative model for how training changes population activity

Although the training-related changes to tuning curves appeared complex in terms of single-cell responses, they could be accurately summarized by a simple quantitative model (Fig. 5).

**Figure 5.**
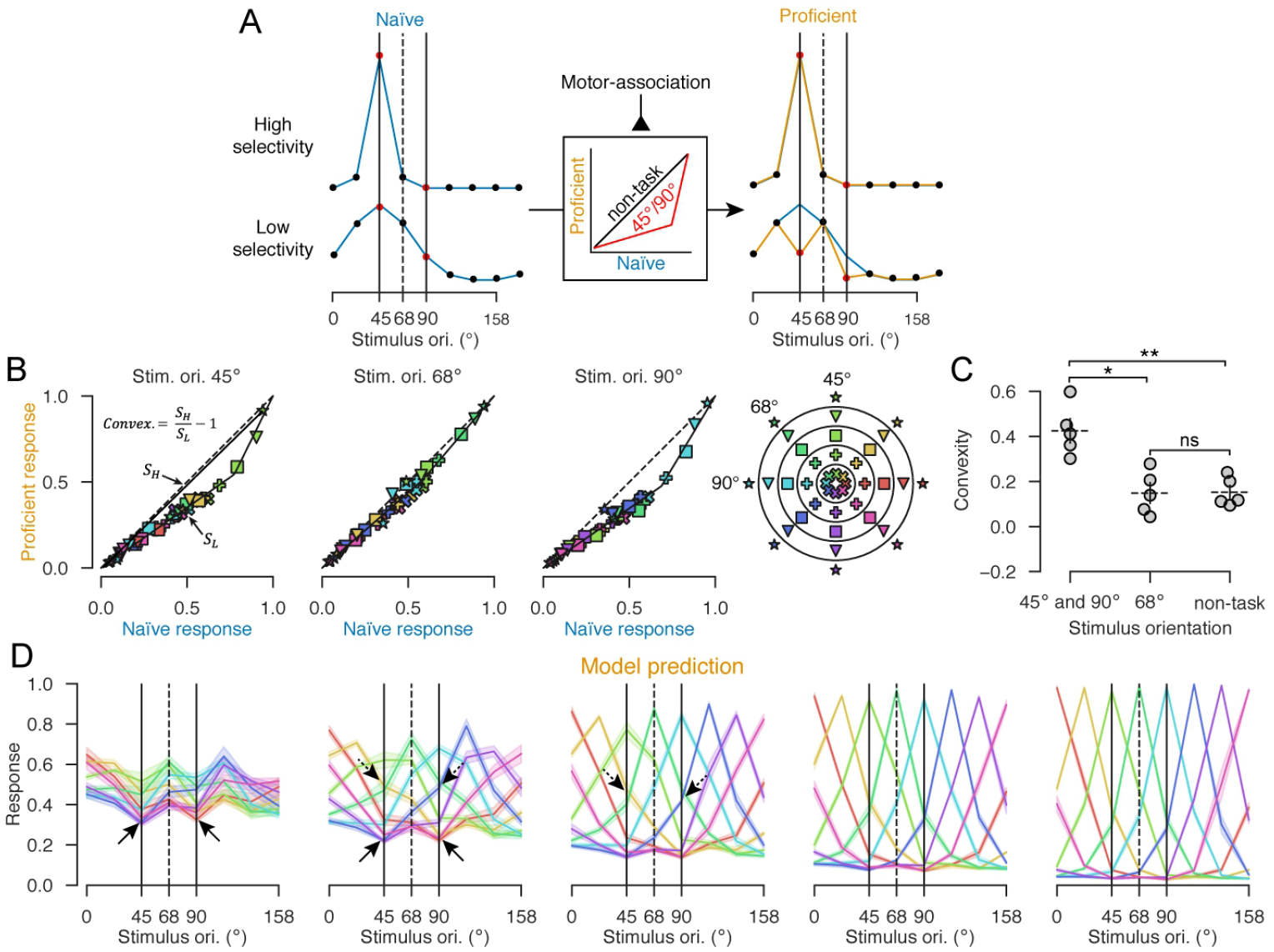
Simple quantitative model for how training changes population activity. **A**, Model schematic. Following task training, the naïve response *f*_*c,θ*_of cell *c* to stimulus *θ* is transformed by nonlinear function *g* _*θ*_, which depends on the stimulus *θ* but not the cell *c*. Blue curves on the left illustrate tuning curves *f*_*c,θ*_of two hypothetical cells in the naïve condition. Middle box illustrates the function *g* _*θ*_, which is more convex for motor-associated stimuli (red curve) than for non-task stimuli (black curve). Orange curves to the right show the proficient responses *g* _*θ*_ (*f*_*c, θ*_), superimposed on original naïve curves (blue). This transformation specifically suppresses moderate responses to the motor-associated stimuli but does not affect strong or zero responses to motor-associated stimuli, or any responses to non-task stimuli. Thus, a cell that was highly selective to 45° is unaffected (top right), while a cell that was weakly selective to 45° develops a multi-peaked tuning curve. **B**, Empirical fits of the function *g*_*θ*_for *θ*= 45°, 68°, and 90°. Each symbol shows the mean response of the same cell groups analyzed in Fig. 2F to the orientation *θ*in naïve vs proficient conditions. Each point shows the average response of cells from all experiments. Black solid lines are stimulus-specific fits of piecewise linear functions *g*_*θ*_relating naïve responses to proficient responses. Symbol color indicates orientation preference and glyph indicates selectivity following the code illustrated in polar coordinates on the right. **C**, Convexity of *g*_*θ*_(defined as the ratio of slopes of the two solid lines in panel B, minus 1; see Methods), for motor associated orientations 45° and 90°, distractor orientation 68°, and all other non-task orientations. Points indicate individual mice. Error bars: mean and SEM (N = 5 mice). Significance stars: paired samples *t*-test. **D**, Proficient orientation tuning curves predicted by the model, obtained by applying the functions fit in **B** to naïve tuning curves. Solid and dashed arrows highlight the same features seen in the actual proficient responses, as shown in Fig. 2F-G. Shading: SEM (N = 5 mice). *, p < 0.05; **, p < 0.01

In this model, visuomotor learning transforms V1 population responses by applying a convex piecewise-linear transformation to the response of each cell (Fig. 5A): if cell *c*’s response to orientation *θ* was *f*_*c, θ*_ before training, then after training it is *f*′_*c, θ*_ = *g*_*θ*_ (*f*_*c, θ*_). Importantly, the function *g*_*θ*_ depends on the stimulus *θ*, but not on the cell *c*: to predict how a cell’s responses to stimulus *θ* transforms after learning it suffices to know the cell’s naïve response to this stimulus *θ* only; other properties of the cell, such as its responses to other stimuli, tuning sharpness, or preferred orientation, provide no additional information. To estimate the functions *g*_*θ*_, we fit piecewise linear functions relating naïve and proficient responses separately for each experiment and orientation, and found that these functions accurately summarized the effects of task training (Fig. 5B). The functions *g*_*θ*_ were most convex for motor-associated stimuli, indicating that cells that responded modestly to these stimuli will have their responses further suppressed after training, but cells that responded either strongly or not at all will be unaffected. For distractor or non-task orientations, the functions were close to linear, indicating little plasticity of responses to these stimuli (Fig. 5C; convexity of motor-associated vs 68°: p = 0.014; motor-associated vs non-task: p = 0.009; 68° vs non-task: p = 0.827, paired samples *t*-test, n = 5 mice). Applying this transformation to the naïve tuning curves, we were able to predict neuronal responses in proficient mice with remarkable accuracy (Fig. 5D; compare to Fig. 4G). Similar results were observed at the level of individual neurons whose activity we tracked between naïve and proficient conditions (Supplementary Fig. 5D). Thus, one simple principle summarizes the representational plasticity we observed: responses to motor-associated stimuli are suppressed by task learning, but only in cells that responded to them at intermediate levels.

The model provides a simple, quantitative explanation for the apparently complex effects of training on single-cell tuning curves changes seen earlier (e.g. Fig. 1I, 4A, 4G). It explains why training affects mostly the cells that are broadly tuned and gives them multipeaked tuning curves: these cells exhibit intermediate levels of response that are affected most by the convex nonlinearity, and thus suppressed specifically to the task orientations. In contrast, strongly tuned cells always fire close to either the minimum or maximum possible, so are unaffected by the nonlinearity. It explains why population sparseness increases most strongly to task stimuli (Fig. 3B), which we proved mathematically will inevitably follow such a convex transformation of firing rates (Appendix 2.1) and will in turn be accompanied by orthogonalization (Appendix 2.2).

This plasticity we observed was better modeled by transformation of population responses than sharpening of orientation tuning curves. To show this we fit a second quantitative model, which transformed each cell’s tuning curve rather than transforming each stimulus’ population response. Specifically, for each cell class *c* (i.e., each orientation preference and selectivity class, corresponding to one curve in Fig. 4F) we fit a piecewise-linear transformation *g*_*c*_ relating activity in naïve and proficient conditions (Supplemental Fig. 6A). This sharpening model did not predict the multi-peaked tuning curves and asymmetric slope shifts observed for weakly tuned neurons following training (Supplemental Fig. 6B) and performed worse on cross-validated metrics of accuracy (Supplemental Fig. 6C), indicating that a general sharpening of individual cell tuning curves insufficiently describes our observed effects.

**Figure 6.**
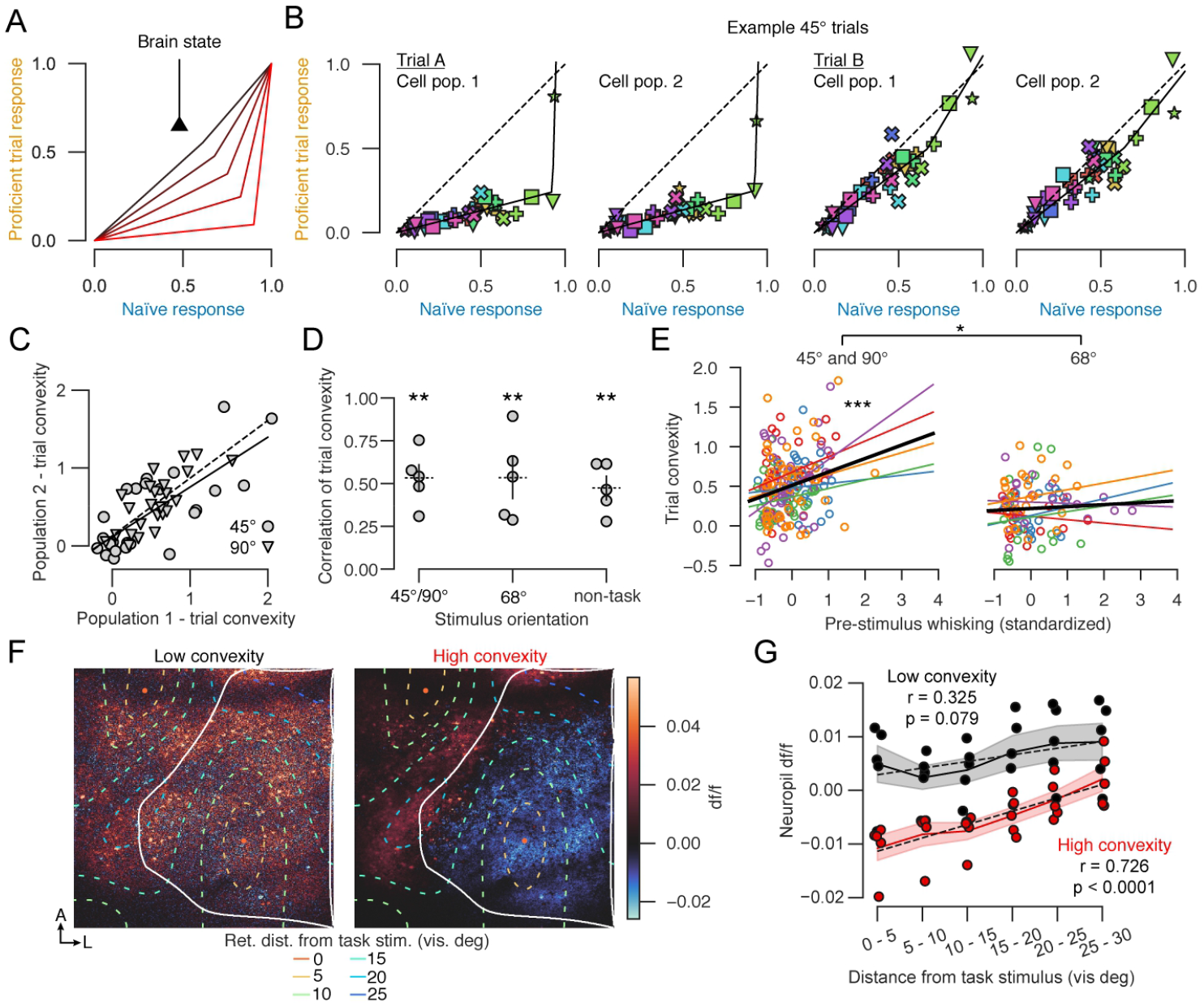
Tuning curve transformation varies dynamically from trial to trial depending on behavioral state. **A**, Dynamic sparsening model: activity undergoes varying levels of sparsening on different trials, depending on instantaneous brain state. **B**, Single-trial transformation functions for two example presentations of 45° gratings in the same recording session, plotted as Fig. 3B. For each trial, responses of separate halves of the cell population are shown. **C**, Similarity of single-trial convexities between two different halves of the cell population, for the recording in **B**. Each point represents a single presentation of the 45° (circle) or 90° stimulus (triangle). Solid and dashed lines are fits to the 45° and 90° trials, respectively. **D**, Correlation of single-trial convexities between two halves of cells (average of 2000 random splits, see Methods), with each point representing average over motor-associated, distractor, or non-task stimuli in one experiment. Error bars: mean and SEM (N = 5 mice). Significance stars: one sample *t*-test for difference to 0. **E**, Correlation of trial convexity with pre-stimulus whisking. Each point represents a stimulus presentation, color coded by mouse identity. Colored lines are linear regression fits for individual mice, black line the mean over mice. Left: motor-associated stimuli; right: distractor stimuli. Significance stars: linear mixed effects model (n = 303 trials, N = 5 mice). **F**, Trial-to-trial variability of neuropil responses. Left and right plots show mean df/f (2 s post-stimulus vs. 1 s pre-stimulus) of two-photon imaging frames to motor-associated orientations. Left and right images show averages over trials for which the population convexity (defined by the ratio of slopes as in panel B) was low (< 0) or high (> 0.3), respectively. Colored contours: retinotopic distances from task stimulus location (see legend). **G**, V1 neuropil responses to motor-associated orientations, as a function of retinotopic distance from the task stimulus, for trials with low and high convexity. Dashed lines: linear regression fits. Shading: SEM (N = 5 mice). *, p < 0.05, **, p < 0.01, ***, p < 0.001.

### Tuning curve transformation varies dynamically from trial to trial depending on behavioral state

Plasticity of cortical representations is often assumed to arise from long-term modifications of local excitatory synapses that changes the sensory drive received by cortical neurons^45,46^. However, the fact that plasticity is better fit as a transformation of single-stimulus population responses than of single-cell tuning curves suggests an alternative hypothesis. Under this hypothesis, V1 neurons receive a visual sensory drive unaffected by training, but motor-associated stimuli engage a circuit process that suppresses the firing of cells receiving weak sensory drive while sparing strongly driven cells. Multiple physiological mechanisms could underlie this process, for example, if motor-associated stimuli caused increased activation of a particular inhibitory cell class, input pathway, or neuromodulatory system. For example, a simple mathematical argument shows that convex transformation of firing rates should be expected from increased subtractive inhibition (Appendix 2.3).

This hypothesis makes an experimental prediction: the convexity of the transformation affecting responses should vary from one trial to the next (Fig. 6A). While a static plasticity mechanism would lead to tuning-curve plasticity that is constant across trials, a dynamic mechanism might be engaged more on some trials than others, due to fluctuations in brain state. Thus, if the degree of transformation in sensory responses varies between stimulus repeats, it indicates a dynamic mechanism. Furthermore, since the circuit process would affect all neurons similarly, trial-to-trial variations in response transformation should be consistent across the population. Finally, the degree of transformation on each trial might correlate with behavioral state. Trial-to-trial variability in neuronal sensory responses is well-documented and has been reported to take additive and multiplicative forms^47–49^. The current hypothesis predicts a further type of trial-to-trial variability in the responses to task-relevant stimuli.

We tested this prediction in the population responses in proficient mice on single trials, and found it to be correct (Fig. 6B-E). We divided cells randomly into two groups, balanced for orientation preference and selectivity, and within each cell group, examined the transformation from trial-averaged population responses in the naïve condition to single-trial population activity in the proficient condition. The convexity of the transformation varied substantially between trials, even within repeats of a single stimulus orientation, but was consistent across the cell groups (Fig. 6B-D; correlation coefficient significantly exceeds 0 at p < 0.05 for each stimulus orientation, one sample *t*-test, n = 5 mice). This consistency in the single-trial estimates obtained from separate cell groups shows that the trial-to-trial variability observed results not from fitting to random noise, but from a consistent population-level transformation of stimulus responses.

The strength of the transformation on a given trial depended on behavioral state. Indeed, the convexity of the transformation depended on whisking prior to stimulus onset and this dependence was specific to the responses to motor-associated stimuli (Fig. 6E, Supplementary Fig. 7; linear mixed effects model: p < 0.0001 for effect of whisking on convexity for motor-associated stimuli; p = 0.014 for difference in strength of this effect between motor-associated and distractor stimuli; p = 0.001 for difference in effect strength between motor-associated and non-task stimuli; n = 303 trials, N = 5 mice). This dependence on behavioral state did not result from stimulus-evoked body movements, as these did not vary between different grating orientations (Supplementary Fig. 2).

Suppression of cortical activity by motor-associated stimuli was strongest in the part of V1 topographically representing the task stimulus (Fig. 6F-G; Supplementary Fig. 8). This could be seen even at the level of neuropil fluorescence, which was suppressed most strongly on trials where the transformation of population activity was most convex, specifically in parts of V1 retinotopically aligned with the stimulus (Fig. 6F-G), and specifically for the motor-associated orientations (Supplementary Fig. 8D). Analysis of individual cells gave consistent results, with cells in cortical areas less than 10° of retinotopic visual angle from the stimulus location showing significant suppression to task orientations, but those greater than 20° not doing so (Supplementary Fig. 8E). These results are thus consistent with a circuit mechanism that suppresses cortical activity, specifically in the region of V1 aligned with the stimulus, with the amount of local suppression varying between trials but strongest when motor-associated stimuli are presented.

## Discussion

We have shown that training in a visuomotor task sparsens and orthogonalizes the population responses of primary visual cortex to oriented stimuli, particularly for orientations associated with actions. These changes could be explained by a simple quantitative law: after training, population responses are transformed by a nonlinear convex function, which is the same for all neurons, but whose convexity varies from trial to trial and is largest on average for motor-associated stimuli. This convex transformation sparsens population responses to motor-associated orientations by suppressing neurons responding at intermediate levels, making the resulting population vectors more orthogonal. This orthogonalization does not increase the fidelity of stimulus coding, which was already perfect: an ideal observer could have perfectly decoded stimulus orientation from the population responses even before training. Rather, orthogonalization might help downstream circuits produce different motor responses to the two task-associated orientations, by biasing even suboptimal decoders to distinguish those task orientations.

The way task training transformed stimulus coding appeared complex if analyzing each cell’s tuning individually but became simple when described at the population level. Single-cell tuning curves showed diverse changes including the development of multimodality and asymmetry. Training transformed the activity of all cells by a nonlinear function that depended on the stimulus but not the cell; an alternative model in which the transformation depended on the cell but not the stimulus could not fit our results. This transformation is much simpler than one might have expected to predict a cell’s response to stimulus following learning, the only thing one needs to know about the cell is its response to that stimulus before learning. No other details of the cell, such as its responses to other stimuli, preferred orientation, or orientation selectivity, are required. Thus, the simplicity of the transformation from naïve to trained responses is only apparent when analyzing all cells together.

This simple population-level effect can explain several of the apparently diverse effects of visuomotor task training previously observed in visual cortex at a single-cell level: it predicts a reduction in the number of cells responding modally to the trained orientations^4,16^, an asymmetrical increase in tuning curve slope specifically at these orientations^13^, and the suppression of neuronal activity following learning in particular for cells with preferred orientation close to task stimuli^3,10^, as well the development of multimodal tuning which we believe to be previously unreported. Some apparent discrepancies with previous studies can be explained by differing definitions. For example, while Ref.^11^ reported recruitment of new neurons selective for task stimuli, this primarily reflected a reduction in neurons responding to multiple stimuli^10^, which is consistent with our observations (we re-analyzed data collected in that study confirmed that population responses recorded there sparsen and orthogonalize also after training; Supplementary Fig. 9). The fact that diverse effects and even apparently contradictory phenomena can be explained by a single equation demonstrates the importance of summarizing data with unambiguous equations, and further suggests these phenomena may arise from a single underlying mechanism, for which we suggest possibilities below.

Despite this concordance with previous results in visual cortex, our findings do not appear fully congruent with results from auditory and somatosensory cortex. In these regions, some studies^50–52^ (but not others^26,53^) report that task training or stimulation of neuromodulatory systems under anesthesia increases the number of electrophysiological recording sites responding modally to the task stimuli. We suggest three non-exclusive reasons for this apparent discrepancy. First, it would be surprising if there were only one mechanism by which cortical representations evolve with experience, and it is reasonable to expect that different mechanisms are employed to a different extent in different cortical regions and tasks. In fact, other studies of associative learning in somatosensory or auditory cortex did observe sparsening^26,53^, suggesting that this mechanism is at least sometimes also employed in non-visual cortices; similar phenomena have also been observed in olfactory bulb^27^. Second, methodological differences may explain at least some of the inconsistencies. Our study (like Refs. ^26,53^) used two-photon imaging to record excitatory cells in superficial layers. Other auditory and somatosensory studies have typically used electrophysiological multi-unit recordings, which are biased toward fast-spiking interneurons, and increased activity of these cells is one possible mechanism by which sparsening of pyramidal cell activity could occur. Third, the expansion of sites responding to task stimuli is a transient phenomenon. After continued training or stimulus exposure, expanded maps can “renormalize” to their original state without compromising behavioral performance^54^; furthermore, induction of map expansion by means other than task training can actually worsen task performance^55^, in particular by increasing the rate of false responses to non-target stimuli^56^. Our task required a long training period, potentially allowing time for map expansion to reverse; it also requires differentially responding to the two stimuli while not responding to the similar distractor stimulus, for which map expansion might impair performance.

We did not observe an increase in the fidelity of orientation coding following training, as we found that stimuli could be decoded from population activity with 100% accuracy, even in naïve mice. This result contrasts with some previous studies^1,8,9,11,12^, for which we offer three possible non-exclusive explanations. First, the visual stimuli we were decoding – high contrast full-screen drifting gratings, with orientations separated by 45° and no superimposed noise – were very distinct. The idea that cortical representations of such distinct stimuli would have such low fidelity that decoding them is difficult, is controversial. Indeed, a recent study found that gratings separated by just 1° could be decoded accurately without any task training^25^. The fact that some previous studies have failed to accurately decode such distinct stimuli from V1 activity does not prove it cannot be done: two-photon microscopy is subject to artifacts such as brain movement and neuropil contamination, which, unless corrected with appropriate software, will introduce noise with correlations of precisely the form that compromise decoding^57,58^. Second, activity in mouse V1 encodes not only visual stimuli but also non-visual features such as ongoing movements^37,38^. This non-visual information may compromise decoder performance, particularly for recordings performed during performance of a behavioral task. Third, the performance obtained by any one decoder represents only a lower bound on the performance of an ideal observer, as decoder performance is sensitive to factors such as training set size and regularization parameters, particularly when decoding from large numbers of cells.

Our results do not determine the mechanisms of training-related sparsening, but they do suffice to formulate a hypothesis: that after training, motor-associated stimuli drive an increase in localized but nonspecific inhibition. It is often assumed that changes in cortical representations arise from plasticity of excitatory inputs onto the cells being recorded. This does not seem a likely mechanism for our results however, as long-term changes to synaptic strengths are presumably static, but the strength of population code transformation we observed varies from trial to trial. Instead, these results are more consistent with increased activity of an inhibitory population in the retinotopic location of the task stimulus, whose response to motor-associated stimuli is strengthened following learning, and whose response to distractor stimuli is strengthened to a lesser extent. Our data do not shed light on the inhibitory class responsible, but previous work in visual and auditory cortex^16,53^ has implicated increased inhibition by somatostatin interneurons in reducing cortical responses following passive stimulus exposure. We observed greater sparsening of population responses to motor-associated than distractor stimuli, indicating that passive exposure cannot be a complete explanation for the plasticity we see, but this does not rule out a similar circuit mechanism underlying both phenomena. For example, perhaps increased spatial attention to motor-associated stimuli accelerates the same visual cortical plasticity processes that occur during passive stimulus presentation, leading to greater sparsening of the code for motor-associated than distractor stimuli.

Our test stimuli were full-field gratings that drove wide regions of visual cortex, but we observed sparsening specifically in the region of V1 retinotopically corresponding to the training stimuli. Sparsening had a promiscuous effect on all excitatory neurons in this region: its effect on all neurons could be explained by the same suppressive firing rate transformation, and suppression was also visible in neuropil fluorescence, which correlates with the summed activity of all local neurons^59^. The strength of suppression fluctuated from trial to trial, and was correlated with pre-stimulus whisking. These observations are all consistent with sparsening being mediated by increased activity of a class of local inhibitory neurons which following training become driven by the task stimuli particularly during alert states, but whose activity is also highly variable between trials regardless of state or stimulus. The output of these neurons would nonspecifically suppress all local excitatory neurons by convex transformation of their firing rates. A simple mathematical argument shows that convex transformation of firing rates should be expected from an increase in subtractive inhibition (Appendix 2.3). Furthermore, previous work shows that local feedback inhibition contributes to V1 orientation tuning^60^; that stimulation of parvalbumin-positive interneurons narrows tuning curves in a manner con-sistent with convex transformation of firing rates via subtractive inhibition, and improves behavioral orientation discrimination^61,62^; and that learning can cause changes in inhibitory activity^63,64^.

The precise plasticity mechanism by which these inhibitory neurons would become driven by the motor-associated stimuli is unclear. One possibility is strengthening of inputs onto these interneurons from local pyramidal cells tuned to motor-associated stimuli. Alternatively, the feedback could arise from long-range excitatory or inhibitory input^16^, or from neuromodulators, which also modulate local inhibitory classes^65–67^. If a long-range input is involved, however, this input would have to specifically target the retinotopic location of the stimulus, in order to explain localized suppression of both cellular and neuropil activity. Future experiments may be able to identify the precise type of inhibitory neurons involved and the plasticity mechanisms that cause them to respond preferentially to motor-associated stimuli, and to test the hypotheses that the resulting sparsening of excitatory population activity correlates with, and contributes to improved behavioral performance.

Regardless of the underlying mechanism, the fact that training-related sparsening orthogonalizes of population responses to the motor-associated stimuli suggests a function for this process. The question of why animals so frequently make “incorrect” choices – i.e. choices leading to suboptimal reward – is one of the biggest puzzles in neuroscience. The “noise hypothesis” provided a superficially attractive explanation to this puzzle: suboptimality results from a hard constraint of neural circuits, and when animals do not obtain high reward, it is because they simply cannot do better. While the noise hypothesis might appear to be supported by recordings of small populations^21–23^, it is ruled out by our own and previous^25^ large-scale population recordings, which show that at least high-contrast grating stimuli can be decoded from naïve cortical activity with 100% accuracy. The question thus becomes even more puzzling: why would an animal that could in principle obtain a high rate of reward, still not do so? We hypothesize that the answer lies in the inductive biases or “priors” that have been shaped by evolution. Gratings are not natural stimuli, and if a mouse ever did encounter one in the wild, it is unlikely that the grating’s orientation would be of behavioral significance. Thus, one might expect mice to have a strong bias towards generalizing a behavioral response learned to one grating orientation to another. This strategy would be beneficial in the wild, where appropriate responses will almost always generalize between stimuli as similar as gratings of different orientations. But in our laboratory task, where different orientations require different responses, this inductive bias of default generalization between gratings leads to suboptimal performance that can only be overcome through extensive training. We hypothesize that sparsening and orthogonalization of cortical representations may contribute to breaking this inductive bias, and speculate that similar procedures might provide a new mechanism to boost the capacity of artificial learning systems.

## Acknowledgments

We thank Charu Reddy for assistance with mouse surgeries, Miles Wells, Hamish Forest, and Laura Funnell for assistance with task training, and Michael Krumin for technical assistance with two-photon calcium imaging. We thank Jasper Poort for sharing data from Poort, et al., 2022.

## Contributions

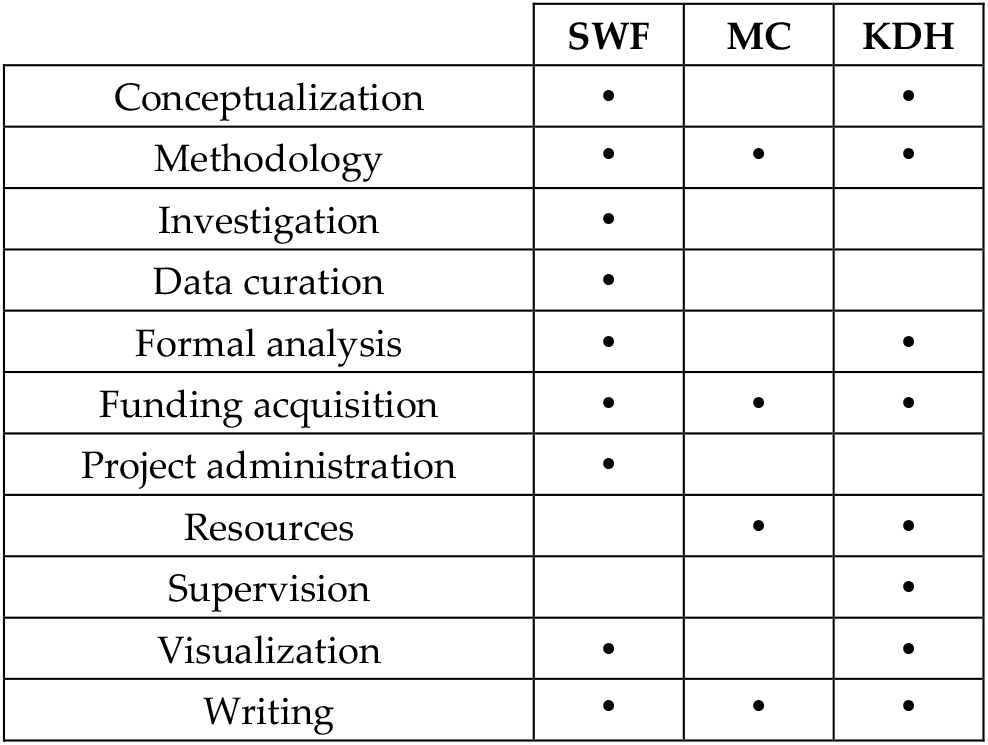

## STAR Methods

### Experimental procedures

All experimental procedures were conducted according to the UK Animals Scientific Procedures Act (1986). Experiments were performed at University College London under personal and project licenses released by the Home Office following appropriate ethics review.

#### Surgical procedure

Five transgenic adult mice (60 days or older) expressing GCaMP6s in excitatory neurons (CaMK2a-tTA;tetO-GCaMP6s) underwent a procedure to implant cortical windows over right primary visual cortex (V1). Mice were anesthetized with isoflurane, an ophthalmic ointment was applied to the eyes, and injections of carprofen and dexamethasone were administered. The hair on the head at the planned incision site was shaved away, and the mouse was transferred to a stereotaxic apparatus where its skull was secured with ear bars. The scalp was cleaned with 70% ethanol to remove loose hairs and other detritus, after which a lidocaine ointment was applied. Following a final application of iodine and ethanol, the scalp over visual cortex was excised, and the edges of the incision were sealed to the skull with a cyanoacrylate adhesive. Using dental acrylic resin, a sterilized metal head plate with a circular well was cemented onto the skull. A 4 mm circular craniotomy was made over right V1 using a biopsy punch, and a glass window was sealed in place with a cyanoacrylate adhesive and dental acrylic resin. At the end of the procedure, mice were removed from anesthesia and placed on a heating pad to recover. Carprofen was added to the mice’s drinking water for three days following surgery to mitigate post-operative pain, and mice were checked daily for any adverse outcomes.

Following recovery, mice were habituated for handling and head-fixation before carrying out recordings.

#### Visuomotor association task

The task is a modification of a two-alternative forced choice contrast discrimination task previously developed by our lab^68^. Mice were head-fixed with their body and hindlimbs resting on a stage, leaving their front forepaws free to turn a small wheel clockwise or anticlockwise. Three computer screens surrounded the mouse, spanning -135 to +135 visual degrees (deg) along the azimuth axis and -35 to +35 v° along the elevation axis. Trials began after 1 - 2 s of continuous quiescence (no wheel movement), after which two full contrast Gabors with sigmas of 18 deg and spatial frequencies of 0.04 cycles/deg were presented simultaneously and centered at -80 and +80 deg azimuth. These Gabors were randomly oriented at either 45°, 68°, or 90°, though the pair were never identical. After an additional quiescence period of approximately 1 s, an auditory cue (12 kHz, 100 ms) would sound, signaling to the mouse that the horizontal position of the Gabors could be manipulated via wheel movement. If the mouse moved the wheel before the auditory cue, the Gabors remained stationary while the quiescence requirement remained in force. When a Gabor was moved to the center screen, a choice was recorded for that trial, and a feedback period was initiated. Correct choices (driving a 45° stimulus to the center, or a 90° stimulus away) were rewarded with 1 - 5 µl of water and a short 0.25 s delay, while incorrect choices (driving a 90° stimulus to the center, or a 45° stimulus away) resulted in a 1 - 2 s burst of white noise. The Gabor was locked at the center position during the feedback, following which it would disappear, and the next pre-trial period of enforced quiescence would begin. During task training, mice were water restricted in line with the approved project license. Mice were considered proficient at the task when they consistently made the correct choice on over 70% of trials.

#### Recording visual responses in V1

Two sessions of two-photon calcium imaging were performed: one before task training (naïve) and one after mice had achieved high performance in the task (proficient). Imaging in the proficient condition was performed immediately after a behavioral session and in the same apparatus.

#### Location of visual areas

Prior to the first two-photon imaging session, we determined the location of V1 in each mouse’s cortical window by recording cortical responses to sparse noise under mesoscopic wide-field calcium imaging and then generating a visual sign map, as previously described ^59^. Mice were placed on a stage of the same type used in the task, and white squares of width 7.5° visual degrees were shown on a black background at a frame rate of 6 Hz for 10 minutes. Squares appeared randomly at fixed positions in a 12 by 36 grid, spanning the retinotopic range of the computer screens. 12% of the squares shown at any one time.

#### Two-photon calcium imaging

Layer 2/3 in V1 was imaged using a commercial two-photon microscope (Bergamo II, Thorlabs Inc) controlled by ScanImage^69^. A Ti:sapphire laser (Chameleon Vision, Coherent) was set to a wavelength between 940 and 980 nm, and the beam was focused with a 16X water-immersion objective (0.8 NA, Nikon). Images were acquired at a frequency of 30 Hz across six planes (5 Hz per plane), a resolution of 512 × 512 pixels, with a frame width between 730 and 810 µm. The fly-back plane was excluded from further analysis. During recordings, mice were head-fixed and placed on the same type of stage used for the task. Three computer screens surrounded the mouse, spanning -135 to +135 v° along the azimuth axis and -35 to +35 v° along the elevation axis.

#### Cell tracking

Neurons were tracked between naïve and proficient recording sessions using a custom-built toolbox written in Python (http://github.com/sfailor/srt4s2p). Mean pixel intensity images, pixel correlation maps, and ROI maps were visualized and recognizable cells and other landmarks were designated in a GUI. The coordinates of these matches were used to perform a perspective transformation to align the naïve and proficient ROI maps. The aligned maps were compared and ROIs whose overlaps met a threshold were manually curated.

#### Sparse noise

To map the retinotopy of V1 under two-photon imaging (Fig. 1C, middle), sparse noise stimuli were presented. Black or white squares of width 4.5° visual degrees were shown on a gray background at a frame rate of 5 Hz for 8 – 30 minutes. Squares appeared randomly at fixed positions in a 16 by 60 grid, spanning the retinotopic range of the computer screens. 1.5% of the squares were shown at any one time.

#### Drifting gratings

At least 16 blocks of drifting grating stimuli were presented in each recording. In each block, gratings spanning 16 directions (22.5° intervals) and a blank stimulus were each presented once in a randomized sequence. Each grating lasted 2 s, with an inter-trial interval sampled randomly from a uniform distribution with a range of 2 – 3 s. Drifting gratings were full contrast and sinusoidal, with a spatial frequency of 0.04 cycles/deg and a temporal frequency of 4 cycles/s, that either encompassed all three screens (fullfield, three mice) or the entire left screen (two mice), contralateral to the recorded hemisphere. Data from the two directions for each of the eight orientations covering 180° were analyzed together.

#### Face recording

An infrared LED illuminated the mouse’s face, and a camera with an infrared filter was used to capture any changes in pupil area or whisking behavior.

### Data analysis

#### Pixel map of retinotopy

To obtain a retinotopic map of the two-photon imaging frame (Fig. 1C middle, Fig. S8A), we analyzed the two-photon recordings during sparse noise stimuli on a pixel-by-pixel basis, without cell detection. Analyses were performed after singular value decomposition (SVD) to accelerate the computation and denoise the data, producing valid results as these computations are linear. First, we z-scored each pixel’s time course independently. Next, we applied single-value decomposition (SVD) on the z-scored image frames, *F* = *USV*^*T*^, where *F* was the full movie encoded as a matrix of size *N*_*pixels*_ × *T, U* was size *N*_*pixels*_ × *N*_*SVDs*_, *S* was a diagonal matrix of singular values, and *V* was size *T* × *N*_*SVDs*_ with *T* being the number of two-photon imaging frames. A matrix *Y* was computed summarizing the mean response of each of the first 100 columns of *V* to each noise frame, as the time-averaged activity in a window 0.2 to 0.6 s after stimulus onset minus the time-averaged activity in a 1 s pre-stimulus window. This matrix was of size *F* × 100, where *F* is the number of noise stimulus frames. The dependence of these responses on individual noise pixels was estimated using ridge regression: *β* = (*X*^*T*^*X* + *λI*)^−1^*X*^*T*^*Y*, where *X* was a *F* × *N*_*noise_squares*_ matrix containing 1 if a particular square was white or black on a particular frame (0 if it was grey), *λ* was a ridge parameter (*λ* = 100), and *I* was the identity matrix. The stimulus dependence of each pixel was then obtained by matrix multiplication *R* = *USβ*, resulting in a matrix *R* of size *N*_*pixels*_ × *N*_*noise_squares*_, encoding the receptive field map of each 2p imaging pixel. To generate retinotopic maps of the imaging frame, each pixel’s receptive field map was smoothed with a Gaussian (sigma 12 v°) and a peak was found, giving a retinotopic position along the elevation and azimuth axes for each pixel.

Pixel retinotopy maps were used to ensure that the two-photon imaging frames were retinotopically aligned with the position of the left task stimulus (0 v° elevation, -80 v° azimuth) during drifting grating recordings. When the optimal imaging location in V1 was identified in naïve mice, an image of the cortical vasculature was saved for positioning subsequent imaging experiments.

#### Visual sign maps

Due to the retinotopic eccentricity of the imaging location in V1 and the large field of view used, it was occasionally the case that areas outside V1 were also recorded. To differentiate V1 from adjacent visual areas, visual sign maps were obtained using the above pixel retinotopy maps averaged across planes (Fig. S8). First, elevation and azimuth maps were smoothed with a median (width 10 pixels) and a Gaussian (sigma 60 pixels) filter. Similar to the process described in Ref. ^70^, the sine of the difference in angle between the gradients of the elevation and azimuth maps was calculated. This sign map was then thresholded to values above 0.31, and pixels that were members of the largest patch were considered to be in V1. This process was consistent in isolating V1, as verified by visual inspection of the elevation and azimuth retinotopic maps.

#### Pixel map of orientation responses

To obtain a pixel map of oriented grating responses (Fig. S8B), the average df/f of each pixel was calculated in response to each stimulus orientation. For each trial, df was defined as the average fluorescence in a post-stimulus window spanning 0 – 2 s, minus the baseline defined as the average fluorescence in a pre-stimulus window spanning -1 to 0 s relative to stimulus onset. This value was divided by f_0_, the baseline measurement. To isolate neuropil responses (Fig. S8D), only pixels that did not belong to a cell, as determined by Suite2P and subsequent manual curation, were included in the analysis.

#### Cell detection

Registration, cell detection, neuropil correction, and deconvolution of the two-photon imaging data were carried out using Suite2P^71^. Imaged planes were aligned with non-rigid registration (four blocks, 128 × 128), and spiking activity was deconvolved from calcium fluorescence using a kernel with a timescale of 2 s.

#### Characterizing single-cell orientation tuning

All cells identified by Suite2P were analyzed for orientation responses. First, each cell’s trial responses were computed by time-averaging its deconvolved activity on each trial over a window of width 0 - 2 s from drifting grating onset. Next, the mean response of each cell to each orientation and to the blank stimulus was computed by averaging over the respective stimulus trials. These mean responses were then normalized by dividing by its mean response to its preferred stimulus condition (either the modal orientation, or mean activity during the blank stimulus if this exceeded mean responses to all orientations).

A cell’s orientation preference was defined in two ways: the orientation it responded maximally to (modal preferred orientation) or its mean preferred orientation, the argument of the complex number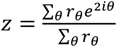, where *r*_*θ*_ is the cell’s mean response to orientation *θ*.The orientation selectivity of a cell was defined as the modulus of *z*. To determine the tuning curve of each cell as a function of its orientation preference and selectivity (Fig. 4F-G), a cross-validated approach was used to avoid erroneously detecting tuning due to random fluctuations in responses. Each cell’s preferred mean orientation and selectivity were calculated using odd-numbered trials, and tuning curves were generated using the mean response to each orientation on even-numbered trials.

Tuning curve slope (Fig. 4H-I) was quantified as the difference between the cell’s response at a stimulus orientation, and the orientation 22.5° closer to the cell’s mean preferred orientation, divided by 22.5. The cell’s tuning curve slope at its preferred mean orientation was defined as the difference between orientations -22.5° or +22.5° from preferred, divided by 45. Thus, if these responses were equal, the tuning curve slope at the preferred orientation would be zero. Tuning curve slopes were separately averaged for all stimulus orientations, and all preferred mean orientations, resulting in nine values per stimulus, per recording. These were displayed as a function of the difference between the stimulus and the mean orientation preference (i.e., -90 to 90 with -90 and 90 being identical). To compare slope magnitudes between stimuli and training conditions, we applied a t-test to the absolute value of the slopes.

#### Discriminability index

The discriminability index (d’) of a cell, its ability to discriminate between two orientations (*θ*_*a*_ and *θ*_*b*_), was defined as 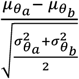 where *μ* and σ^2^ are the mean and variance of the respective orientation responses. The mean and variance for each stimulus orientation was the average of the mean and variance of the two corresponding stimulus directions.

#### Population sparseness

Population sparseness was calculated using the Treves-Rolls formula^39^,

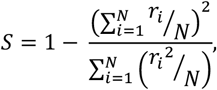

where ***r***_***i***_ is the response of neuron ***i***, and ***N*** is the number of neurons. The Treves-Rolls measure can be thought of as counting the fraction of cells with near-zero activity. For example, if M of N cells respond with equal rate, and the remaining N-M cells are silent, this measure takes the value 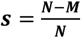.

### Orthogonalization of population responses

To calculate the orthogonalization of population responses between different stimulus orientations (Fig. 3D, S3D), we split the trials into odd and even halves, and computed the *N*_*cells*_-dimensional population response vectors *P*_*i*_(*θ*) to orientation 0 for the trial set *i* (*i* = 1: odd trials; *i* = 2: even trials). We computed the cosine similarity between orientations *θ*_1_ and *θ*_2_ as 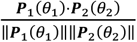. This process resulted in an eight-by-eight matrix of similarity values for each mouse and training condition. Computing this similarity between two separate halves ensured that the diagonal was not 1 by definition.

#### Dimensionality reduction

To illustrate the decodability of population responses in a 2-dimensional plot (Fig. S3A), we trained a linear regression model to decode a 2-dimensional vector (cos 2*θ*, sin 2*θ*) for each trial from the *N*_*ceIIs*_-dimensional population response vector on that trial, where *θ* is the stimulus orientation ranging from 0 to 180°. The model was trained on odd trials, and then applied to population responses on even trials to obtain a two-dimensional projection of population activity that separates points by stimulus orientation.

To visualize the orthogonalization of the 45° and 90° population responses (Fig. 3C), we split the responses to 45° and 90° into odd- and even-numbered trials. Projections were determined by singular value decomposition of the odd-numbered trials, and the even-numbered trials were projected onto the resulting top two components.

#### Stimulus prediction

Orientation was also decoded from population activity using linear discriminant analysis (LDA; Fig. 2B-C, S3B). An LDA model was fit using the population responses in odd trials, and its performance was assessed on even trials. To build the model, we used the class *LinearDiscriminantAnalysi*s from the Python library scikit-learn, with solver set to “eigen” and the shrinkage coefficient automatically calculated.

#### Modeling visuomotor association-evoked changes to orientation responses

For each mouse, cells in the naïve and proficient recordings were divided into classes by binning mean orientation preference (eight bins, *0°:* 168.75 – 11.25°, *23°*: 11.25 – 33.75°, *45°*: 33.75 – 56.25°, *68°*: 56.25 – 78.75°, *90°*: 78.75 – 101.25°, *113°*: 101.25 – 123.75°, *135°*: 123.75 – 146.25°, *158°*: 146.25 – 168.75°) and selectivity (five bins, 0 – 0.16, 0.16 – 0.32, 0.32 – 0.48, 0.48 – 0.64, 0.64 – 1). The mean response of each cell class to each stimulus was determined by cross-validation, using odd trials to determine the cell’s tuning class, and using even trials to compute its tuning, as described above.

Responses in the proficient mice were fit by piecewise linear functions of responses in naïve mice (Fig. 5B). The functions were constrained to pass through (0,0), (*a, b*), and (1,1), where *a* and *b* are parameters. This was achieved using the function *r*_*p*_ = *f*_*a,b*_(*r*_*n*_):

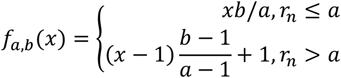

The parameters *a* and *b* were fit for each mouse and stimulus by nonlinear least squares (Python library SciPy, *optimize*.*curve_fit*), constrained to values between 0 and 1.

The convexity of the transformation from naïve to proficient population responses to a stimulus 0 was quantified as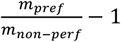, where *m*_*pref*_ was the slope of a line from the origin to the point representing the cell class of preferred orientation *θ* and maximal selectivity, and *m*_*non-pref*_ was the slope of a linear regression on the points corresponding to all cell classes of orientation preference other than *θ*. This approach was used to measure convexity on mean responses, relating the trial-averaged population response in the same mouse prior and after training (Fig. 5B-C), and on single trials, where the population responses in single trial in a proficient mouse was compared to the trial-averaged population response in that mouse prior to training (Fig. 6B).

To assess the consistency of trial-to-trial fluctuations in sparsening across the population (Fig. 6C-D), we randomly divided the proficient cells into two populations balanced for orientation preference and selectivity. Trial-by-trial convexity was measured, as described above, for each cell population, and the correlation coefficient of these convexities was computed. This process was repeated 2000 times, and the average correlation in convexity over orientations was found for each mouse. This approach of independently computing the convexity from two non-overlapping cell populations ensured that the measured trial-to-trial variability did not arise from noise or overfitting, which would be independent between the two groups, but instead reflected genuine and consistent transformations of population activity.

#### Pupil area and whisking

Facial video recordings were processed with the toolkit FaceMap (www.github.com/MouseLand/FaceMap) to obtain traces of pupil area and whisking intensity. The pupil area was defined as the area of a Gaussian fit on thresholded pupil frames, where pixels outside the pupil were set to zero. Whisking intensity was defined as the average change in individual pixels between frames for a region of interest limited to the whisker pad. From these resulting traces, trial-evoked changes in pupil area and whisking were calculated. First, traces of pupil area and whisking intensity were z-scored for each recording session. Second, for each trial pupil area and whisking were averaged in a post-stimulus time window spanning 0 to 2 s. Lastly, stimulus-evoked changes in pupil area and whisking were calculated by subtracting a pre-stimulus baseline, defined as the average pupil area and whisking in a -1 to 0 s window.

#### Statistics

Simultaneously recorded neurons are not statistically independent, so analyses of them must use one of two approaches.

The first approach is to treat the experiment, rather than the neuron, as the unit of independent variability: to summarize the activity of all cells in an experiment in one number, and to use a traditional method such as a t-test, with N being the number of experiments rather than the number of neurons. We did this for the analyses in figures: 2C-E, 2I, 3C, 4D, 4G, 5B, 5D, 6B, S2B, S2D, S3C, S4E, S6E, S8C-D, A1H-K.

The second approach is to use hierarchical models, such as linear mixed effects models, which account for correlated differences in all cells of one recording. We did this for the analyses in figures: 2B, 4E, 5E, S4B, S5A-D, A1A-D, using the following models: 2B: change in response ∼ orientation (1 + mouse); 4E: trial convexity ∼ whisking * stimulus (1 + mouse); 5E: cosine similarity ∼ population sparseness + condition (1 + mouse | condition); S4B: stable ∼ stimulus + (1 + mouse) where stimulus was composed of the stimuli being compared within a selectivity bin; S5A-D: trial convexity ∼ behavior * stimulus (1 + mouse); A1A-D: mean response/coefficient of variation/response index/d-prime ∼ condition + (1 + mouse | condition).

---

**Supplementary Figure 1.**
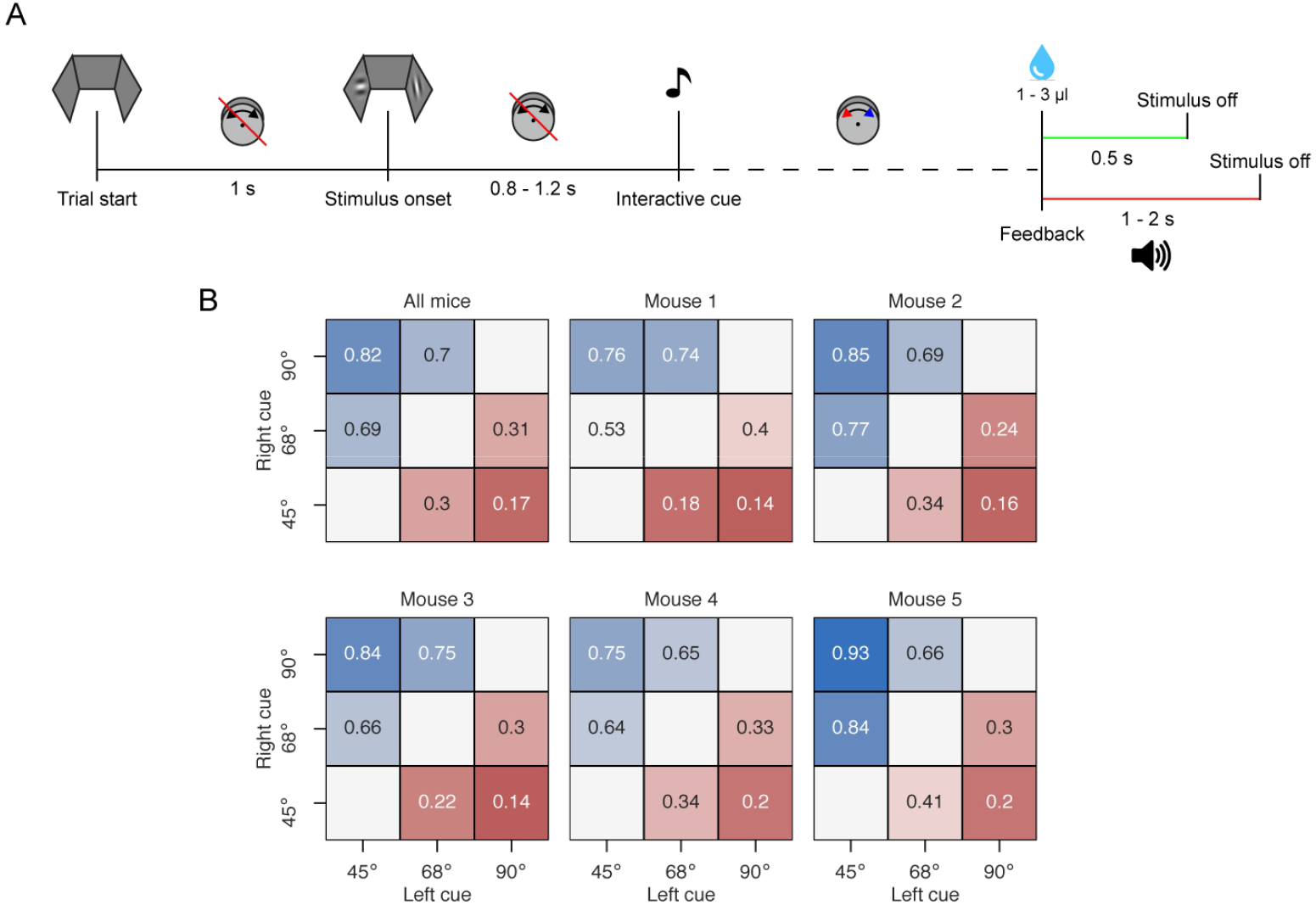
Task details. **A**, Temporal structure of the task. **B**, Behavioral performance for all mice. Matrices show the proportion of clockwise choices for all cue pairings averaged over ten highest performing sessions. Cue pairings that were not presented are shown in white.

**Supplementary Figure 2.**
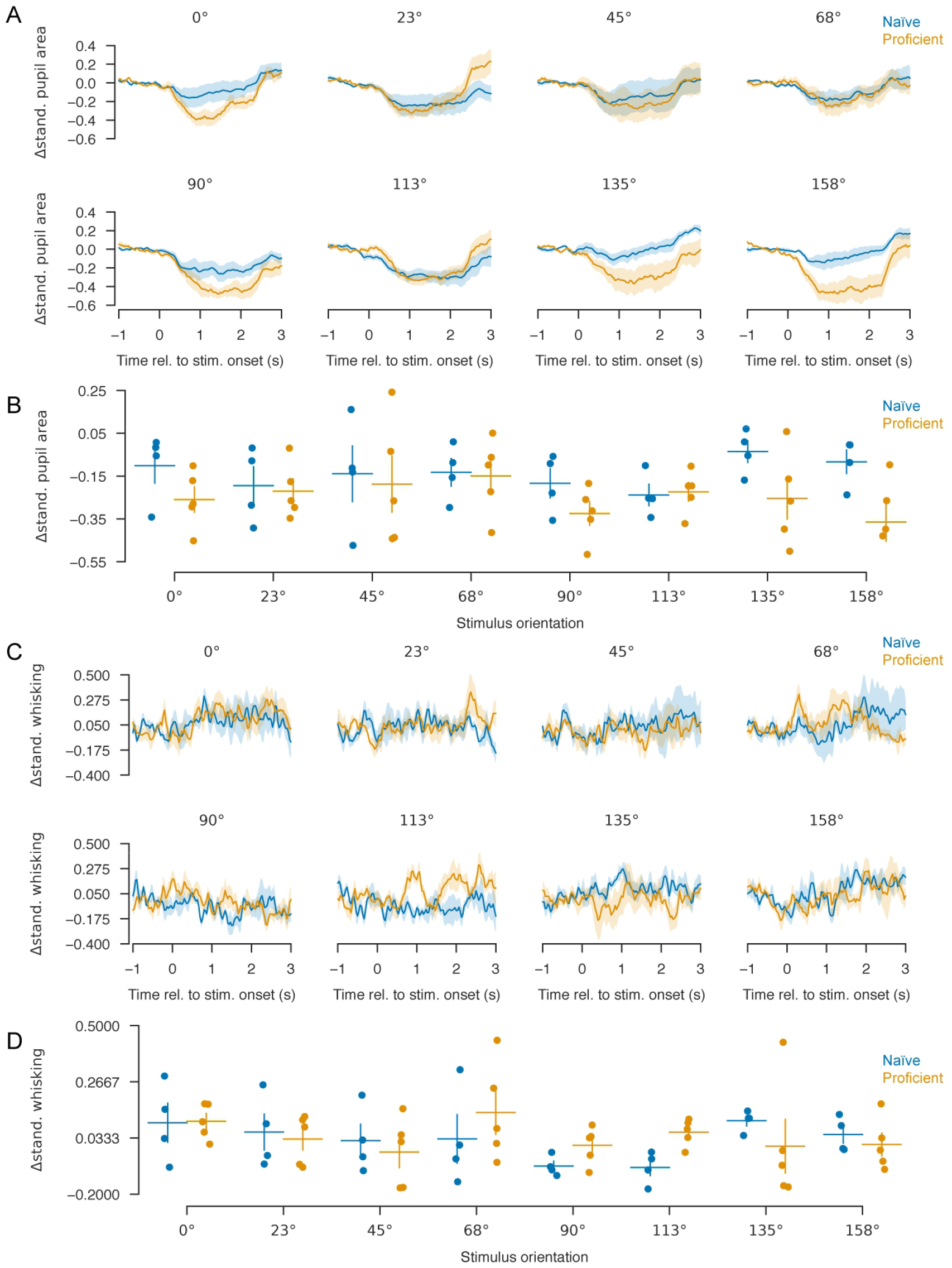
Measures of behavioral responses during passive viewing of grating stimuli. **A**, Stimulus-triggered pupil area time course, averaged over all trials of each stimulus orientation and training condition. Stimulus presentation causes pupil constriction, but pupil responses to motor-associated orientations do not appear substantially different to those to other stimuli. Shaded regions: SEM (n = 5 mice). **B**, Average change in pupil area within gray shaded time windows shown in panel A. ANOVA indicated a marginal effect of training (p = 0.053), and no effect of stimulus orientation (p = 0.279) or their interaction (p = 0.951). Error bars: mean and SEM (n = 5 proficient mice, n = 4 naïve mice, one of which did not have video recorded). (**C** and **D**) Same as in A and B but for whisking, assessed by video motion energy over the whisker pad. ANOVA indicated no significant effect of training (p = 0.547), stimulus orientation (p = 0.061), or their interaction (p = 0.372).

**Supplementary Figure 3.**
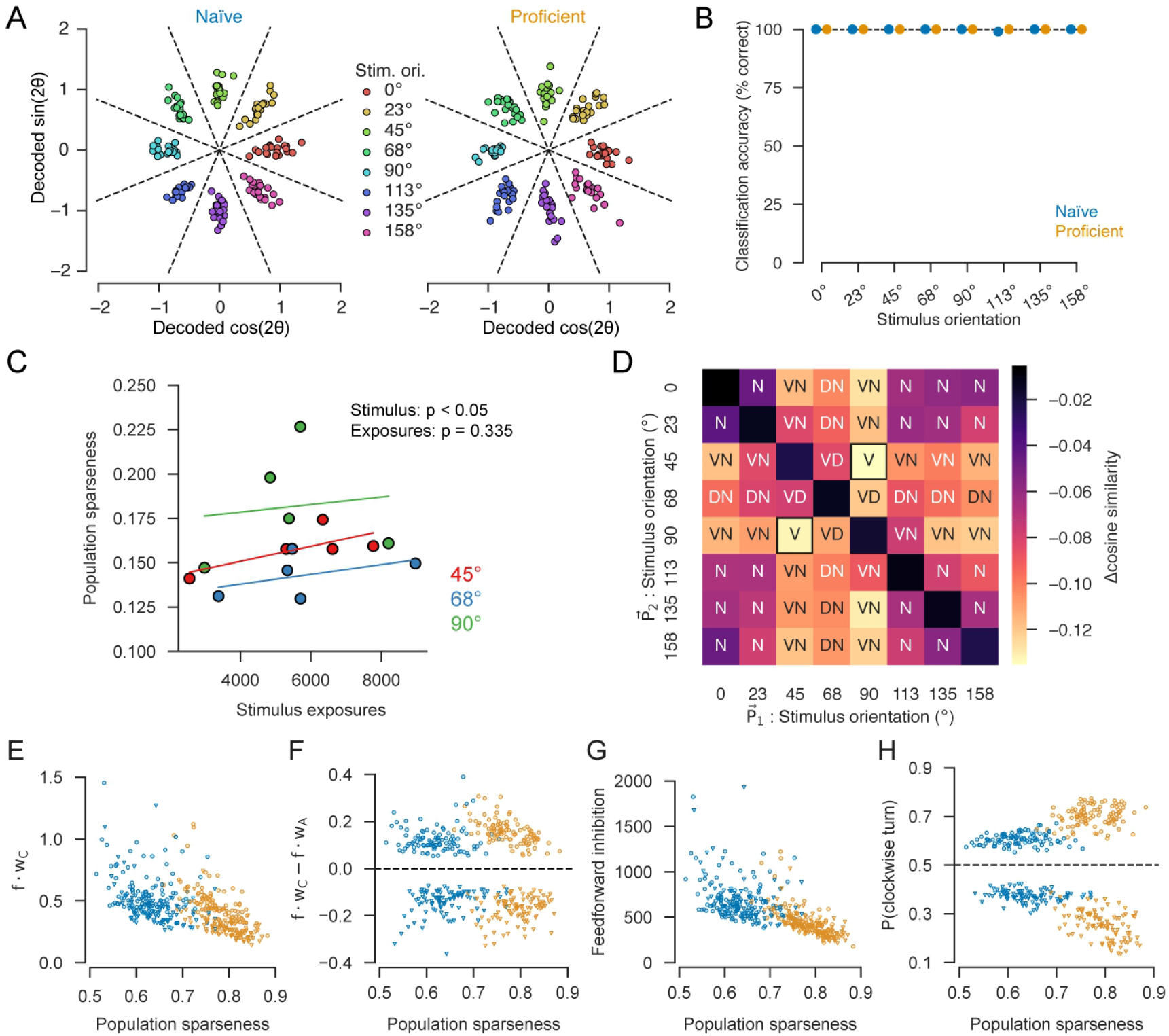
Population-level analyses of task training effects. **A**, Dimensionality reduction indicates perfect separation of population response vectors to different stimuli from one mouse before (left) and after training (right). Each point represents a single-trial response to orientation indicated by the color. X- and y coordinates of each point are weighted sums of neural activity, with weights determined by cross-validated linear regression decoding of *cos*(2 *θ*) and *sin*(2 *θ*) where *θ*is stimulus orientation. **B**, Cross-validated classification accuracy for decoding stimulus orientation from naïve and proficient mice with linear discriminant analysis (here all orientations are decoded, unlike Fig. 2B in which only two are). Dashed line indicates perfect performance (n = 5 mice); shading for standard error is present but too small to see. **C**, Relationship between stimulus orientation and cumulative stimulus exposure on population sparseness. Each point represents one stimulus in one mouse (N = 5 mice, three stimuli per mouse). The x-axis represents cumulative stimulus exposures from the last 20 training sessions. Lines are least squares fits for each stimulus. ANCOVA shows significant effect for stimulus orientation on population sparseness, but not cumulative exposures or its interaction with stimulus orientation (stimulus, p = 0.0332; exposures, p = 0.335). **D**, Heatmap showing change in cosine similarity between mean population responses to each pair of orientations following task training. Annotations correspond to stimulus comparisons used in Fig. 3D. **E-F**, Diagnostics from decision circuit model in Fig. 3F. **E**, Excitatory input to one decision unit, as a function of population sparseness. Each point represents one trial. Blue: naïve condition, orange: proficient. Circles: 45° stimulus, triangles: 90° stimulus. **F**, Difference in excitatory input to the two units. **G**, Feedforward inhibition strength as a function of population sparseness. **H**, Probability of a clockwise turn, as a function of population sparseness.

**Supplementary Figure 4.**
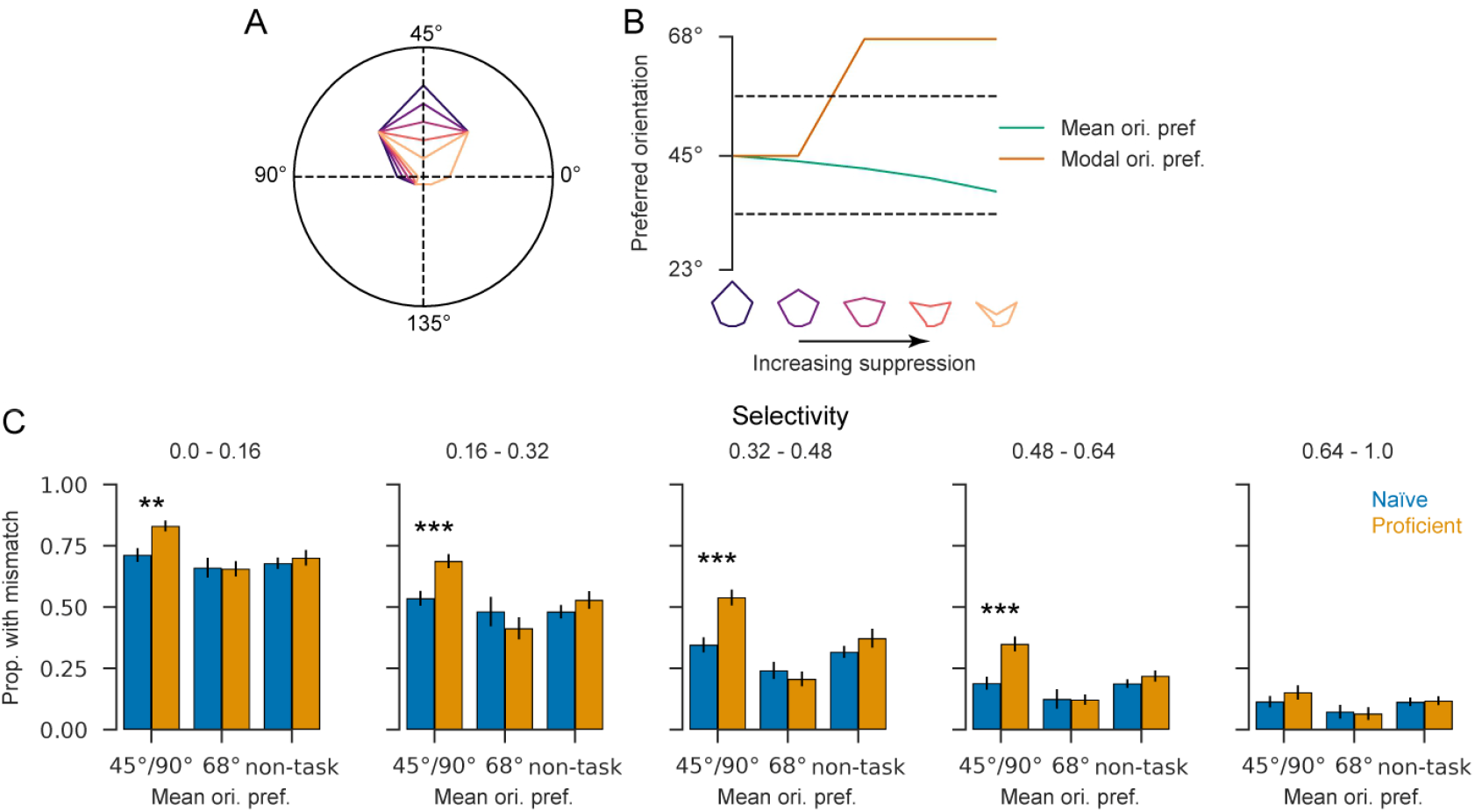
Multimodal tuning curves lead to mismatched mean and modal orientations. **A**, Polar tuning curves for a simulated neuron that is originally symmetrically and unimodally tuned to 45°(purple) undergoing increasing levels of suppression (0%, 20%, 40%, 60%, 80%) at motor-associated orientations 45° and 90° (shades leading to orange). **B**, Circular mean and modal orientation preference of this simulated cell, as a function of suppression level. Increasing the suppression of the response to 45° and 90° results in a stepwise change in modal orientation preference but only a modest change in mean orientation preference. Bottom: polar tuning curve shapes for each suppression level. **C**, Proportion of experimentally recorded neurons with mismatched mean and modal orientations, as a function of training condition (color), mean orientation preference (x-axis), and selectivity (panel). While the modal preference is always one of the stimulus orientations shown, the mean orientation can take any value. We therefore detected a match if the mean orientation was closer to the modal preference than to any other of the stimuli presented; dashed lines in (B) indicate the range of mean orientations that would be matched to a mode of 45°. The fraction of mismatched neurons was significantly larger in trained mice, specifically for neurons with a mean of 45° and 90°, for all selectivity levels except the highest (p = 0.001, p < 0.0001, p < 0.0001, p < 0.0001, p = 0.06 for the 5 selectivity levels, 2-samples t test), but was unchanged for 68° (p = 0.90, p = 0.36, p = 0.32, p = 0.96, p = 0.85) and non-task (p = 0.35, p = 0.07, p = 0.10, p = 0.13, p = 0.74, paired samples *t-*test) orientation preferences. Error bars: SEM (N = 5 mice). **, p < 0.01; ***, p < 0.001.

**Supplementary Figure 5.**
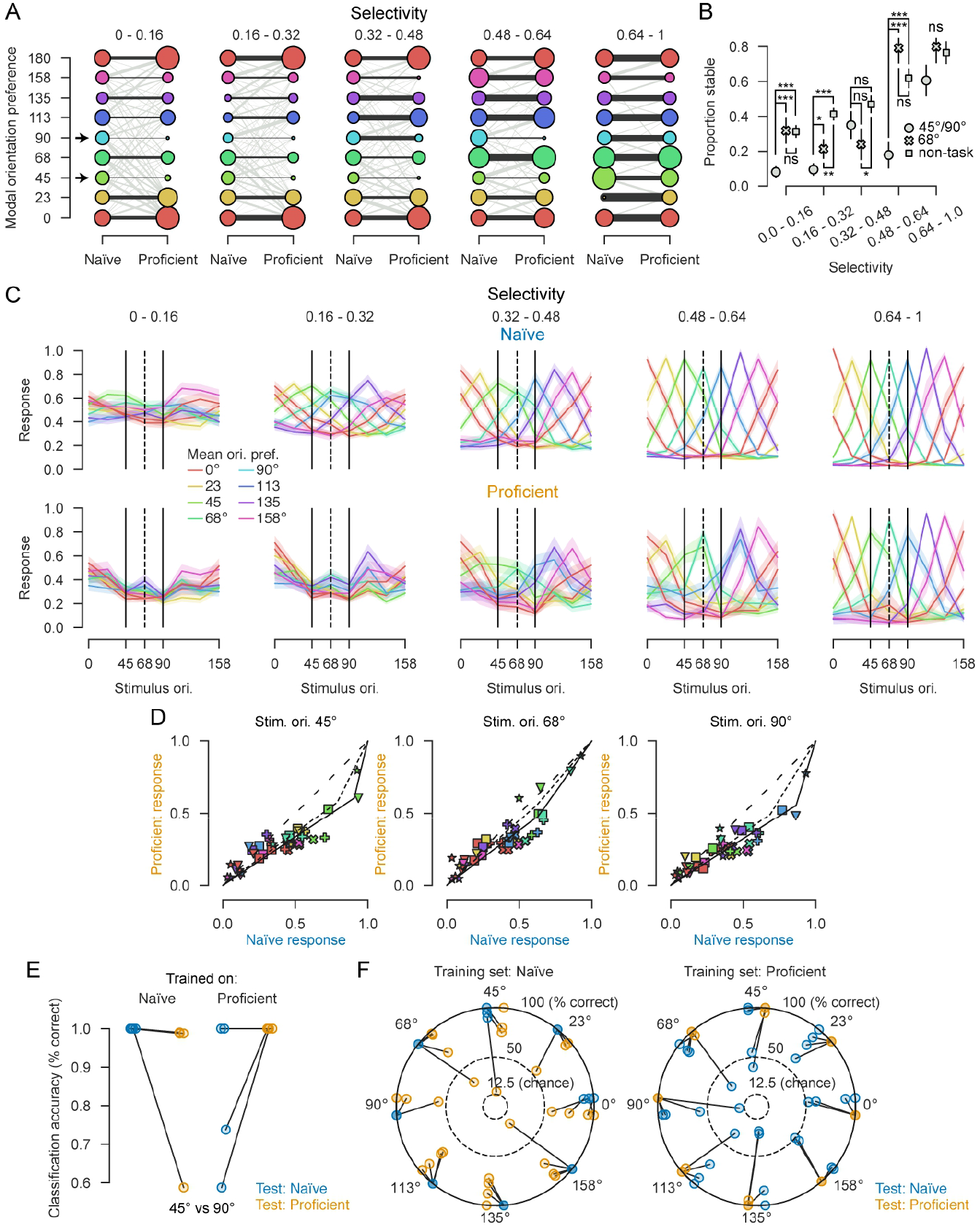
Changes in tuning curves of neurons tracked across training. **A**, Transition plot showing the change in modal orientation preference of neurons tracked from the naïve to the proficient condition. Neurons were divided into five groups according to orientation selectivity in the naïve condition (panels). Each cell’s modal orientation preference was computed in naïve and proficient conditions. The diameter of the circles in each vertical position represents the proportion of neurons with each modal preference, with left and right columns for naïve and proficient conditions. Red circles corresponding to 0° are replicated at the top as 180° to avoid imposing an artificial discontinuity. The thickness of the line connecting two circles indicates the fraction of neurons from each naïve modal preference which transition to each proficient model preference, colored light grey for cells whose preference changed and black for cells of stable orientation preference. Arrows mark motor-associated orientations 45° and 90°; note the small fraction of stable cells for these orientations, particularly for neurons of low naïve selectivity. **B**, Proportion of neurons with stable modal orientation preferences, as a function of naïve orientation preference and tuning strength. Cells that were weakly tuned to motor-associated orientations in naïve conditions had significantly less stable modal preference than those preferring distractor or non-task orientations. Selectivity 0 to 0.16, 45° and 90° vs 68°: p < 0.0001. 45° and 90° vs non-task: p < 0.0001. 68° vs non-task: p = 0.947. Selectivity 0.16 to 0.32, 45° and 90° vs 68°: p = 0.04. 45° and 90° vs non-task: p < 0.0001. 68° vs non-task: p = 0.001. Selectivity 0.32 to 0.48, 45° and 90° vs 68°: p = 0.287. 45° and 90° vs non-task: p = 0.216. 68° vs non-task: p = 0.03. Selectivity 0.48 to 0.64, 45° and 90° vs 68°: p < 0.0001. 45° and 90° vs non-task: p < 0.0001. 68° vs non-task: p = 0.157. Selectivity 0.64 to 1, 45° and 90° vs 68°: p = 0.097. 45° and 90° vs non-task: p = 0.087. 68° vs non-task: p = 0.877. Error bars: mean and SEM; significance stars: hierarchical linear mixed effects model (n = 1107 cells, N = 4 mice). *, p < 0.05; **, p < 0.01, ***, p < 0.001. **C**, Average orientation tuning curves for cell groups tracked across training conditions, defined by mean orientation preference (color) and selectivity (column) in naïve mice. This plot is made similarly to Fig. 4F-G, except that here the proficient tuning curves are averages over cells grouped by their naïve tuning preference, which is only possible for tracked cells. Despite this difference, the tuning curves appear very similar to those in Fig. 4. This excludes the possibility that changes in the average tuning of cells grouped by preferred orientation and tuning strength were due to cells moving between these groups, rather than changes in the tuning of the cells. **D**, Empirical fits of the function *g*_*θ*_for *θ*= 45°, 68°, and 90°, as done in Fig. 5B, for neurons tracked across training and grouped by their naïve preferences (solid line). Dashed line is the fit to data from all neurons, as shown in Fig. 5B. Due to the smaller number of tracked cells, convexity could only be estimated from the cells of all experiments combined, thus statistical analysis of convexity could not be performed for tracked cells alone. **E**, Accuracy of linear discriminant binary prediction of 45° vs 90° from activity of tracked cells only, training on either naïve or proficient data, and testing on held-out naïve or proficient data. **F**, Same analysis for all orientations, plotted in polar coordinates with radius representing classification accuracy and angle representing the stimulus being decoded (the slight angular shift between blue and orange points is for visualization purposes only). Although a significant difference in accuracy was not found between any test-set groups (p > 0.05, paired-samples *t*-test), this likely reflects the inherent conservatism of t-tests in conditions of high variance: accuracy trended worse when the decoder was tested on a condition different to what it was trained on.

**Supplementary Figure 6.**
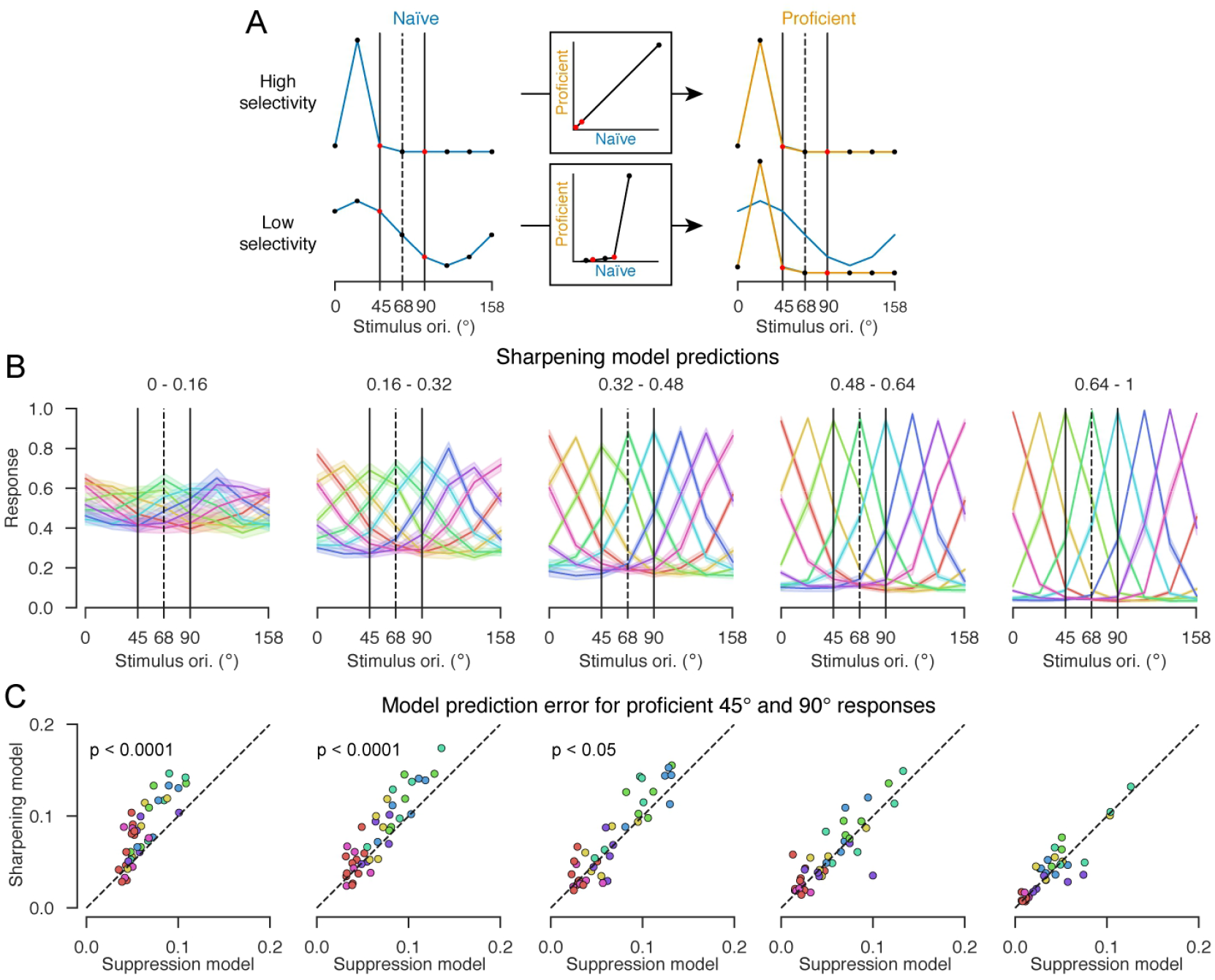
Cell-specific sharpening model does not explain training-evoked effects on tuning curves. **A**, Model schematic. Following task training, the naïve response *f*_*c,θ*_ of cell *c* to stimuli *θ*is transformed by nonlinear function *g*_*c*_, which is specific to the cell *c*, but common across stimuli (in contrast to the model of Fig. 5, which is specific to the stimulus not the cell). Blue curves on the left illustrate tuning curves *f*_*c,θ*_ of two hypothetical cells in the naïve condition. Middle box illustrates two functions *g*_*c*_, which are fit separately to maximize the fit for each cell group. Right: orange curves show the proficient responses *g*_*c*_(*f*_*c,θ*_), superimposed on original naïve curves (blue); red dots are the motor-associated stimuli 45° and 90°. For the neuron with lower selectivity (bottom), training sharpens the tuning curve. **B**, Predicted tuning curves from the optimal fit of the sharpening model, which cannot recapitulate the asymmetric and multipeaked tuning curves observed following training (Fig. 4G). **C**, The stimulus-specific suppression model illustrated in Fig. 5A is quantitatively superior at capturing the suppression of 45° and 90° for neurons with low selectivity in the proficient condition, as demonstrated by lower cross-validated prediction error (paired Student’s *t*-test, n = 5 mice, 40 cell types per mouse).

**Supplementary Figure 7.**
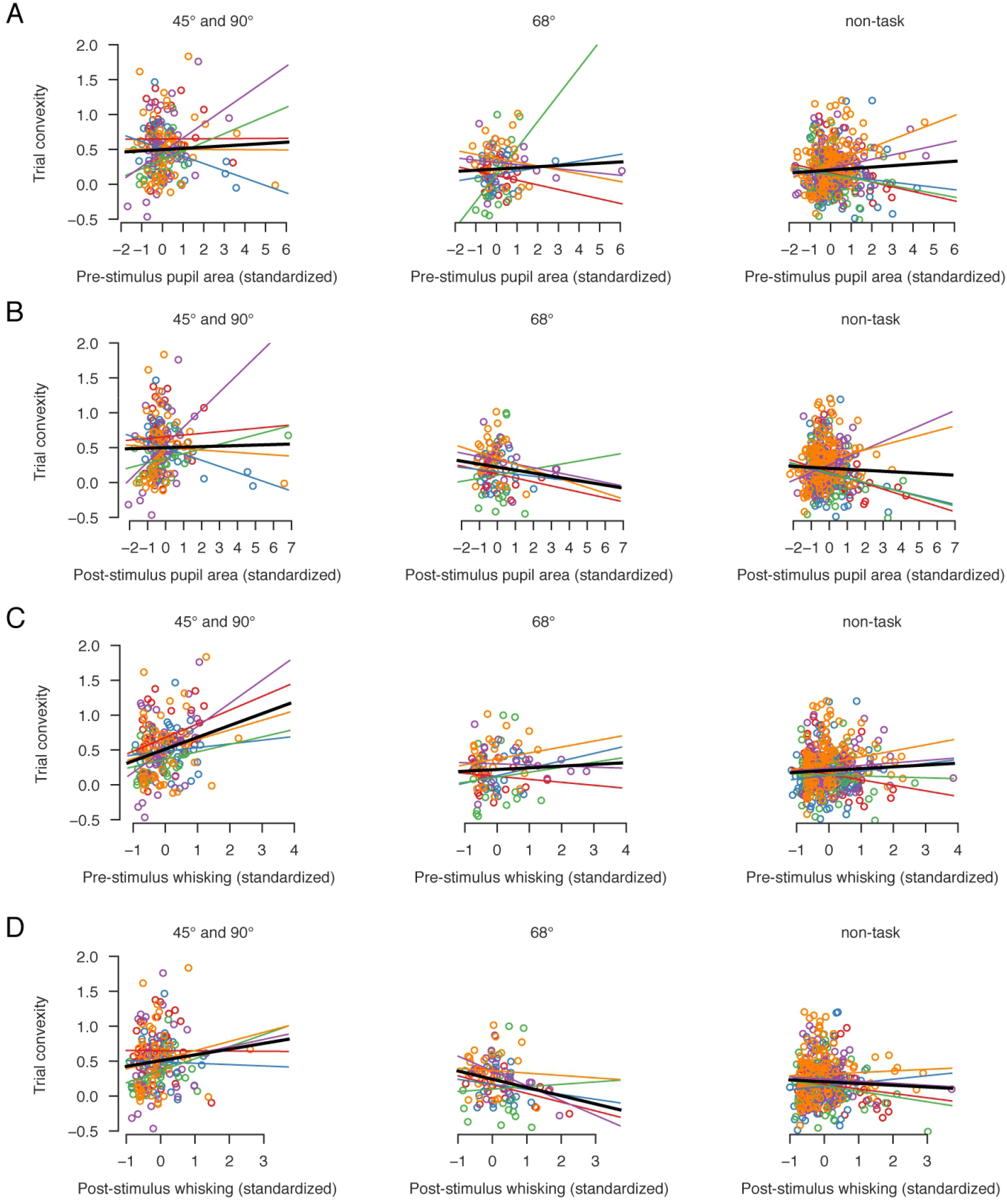
Correlation of trial convexity with multiple measures of behavioral state. **A**, Correlation of single trial convexity pre-stimulus pupil area, plotted as in Fig. 6E, for motor-associated stimuli (left), distractor stimuli (center), and non-task stimuli (right). **B-D**, similar plots for post-stimulus pupil area, pre- and post-stimulus whisking. Convexity correlated positively with pre-stimulus whisking for the motor-associated stimuli (linear mixed effects model; p <0.0001) and the effect was significantly larger than for distractor (p = 0.014) and non-task stimuli (p = 0.001). Convexity was not significantly correlated with post-stimulus whisking for the motor-associated stimuli (p = 0.065), but the distractor (p = 0.006) and non-task stimuli (p = 0.028) showed significant lower levels of correlation. The correlation of the distractor stimulus with post-stimulus whisking was significantly negative (linear mixed effects model; p = 0.009).

**Supplementary Figure 8.**
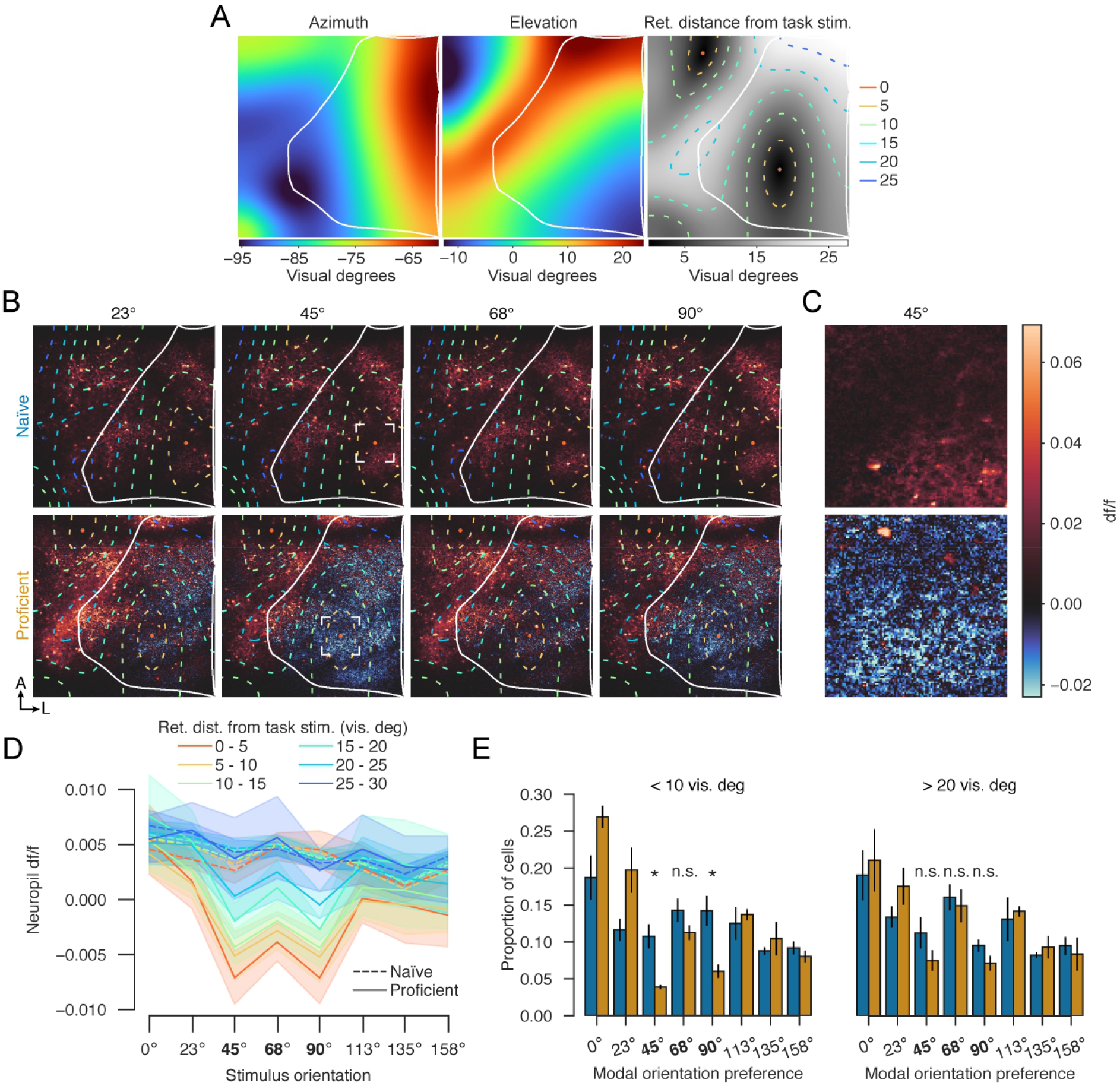
Response suppression is aligned with the retinotopic location of the task stimulus. **A**, Retinotopic mapping of visual cortex, for an example mouse. Left two pseudocolor plots show preferred azimuth and elevation for each pixel in the field of view, assessed by analyzing responses to sparse noise stimuli. White line demarcates the border of V1. Right panel shows distance in degrees of visual angle from each pixel’s preferred retinotopic location to the retinotopic position of the task stimulus, in pseudocolor (gray-scale), and with contour representation (dashed colored lines). **B**, Mean df/f of two-photon imaging frames during presentation of full-field gratings of the marked orientations in the same mouse prior to (top) and after training (bottom). White lines and colored contours mark V1 boundary and retinotopic distance to stimulus location, as in **A. C**, Zoom into boxed regions of B (45° orientation). Note that after training, neuropil is suppressed in the region retinotopically matching the stimulus, although individual cells continue to respond strongly there. **D**, V1 neuropil responses as a function of stimulus orientation and retinotopic distance from the task stimulus position (colors), for naïve and proficient mice (dashed and solid lines). Shading: SEM (n = 5 mice). Note specific suppression of responses to task orientations in pixels retinotopically close to the stimulus location. **E**, Histogram of modal orientation preferences of V1 cells in naïve and proficient mice, for cells close to (left) and distant from (right) the retinotopic position of the task stimulus, plotted as in Fig. 4D. The proportion of cells preferring 45° and 90° but not 68° changes significantly amongst cells within 10 v° of the task stimulus location (p = 0.020, p = 0.045, p = 0.121, paired samples *t*-test). For cells further than 20 v° from the task stimulus location, all three changes are insignificant (p = 0.206, p = 0.132, p = 0.762, paired samples *t*-test). Error bars: SEM (n = 5 mice). *, p < 0.05.

**Supplementary Figure 9.**
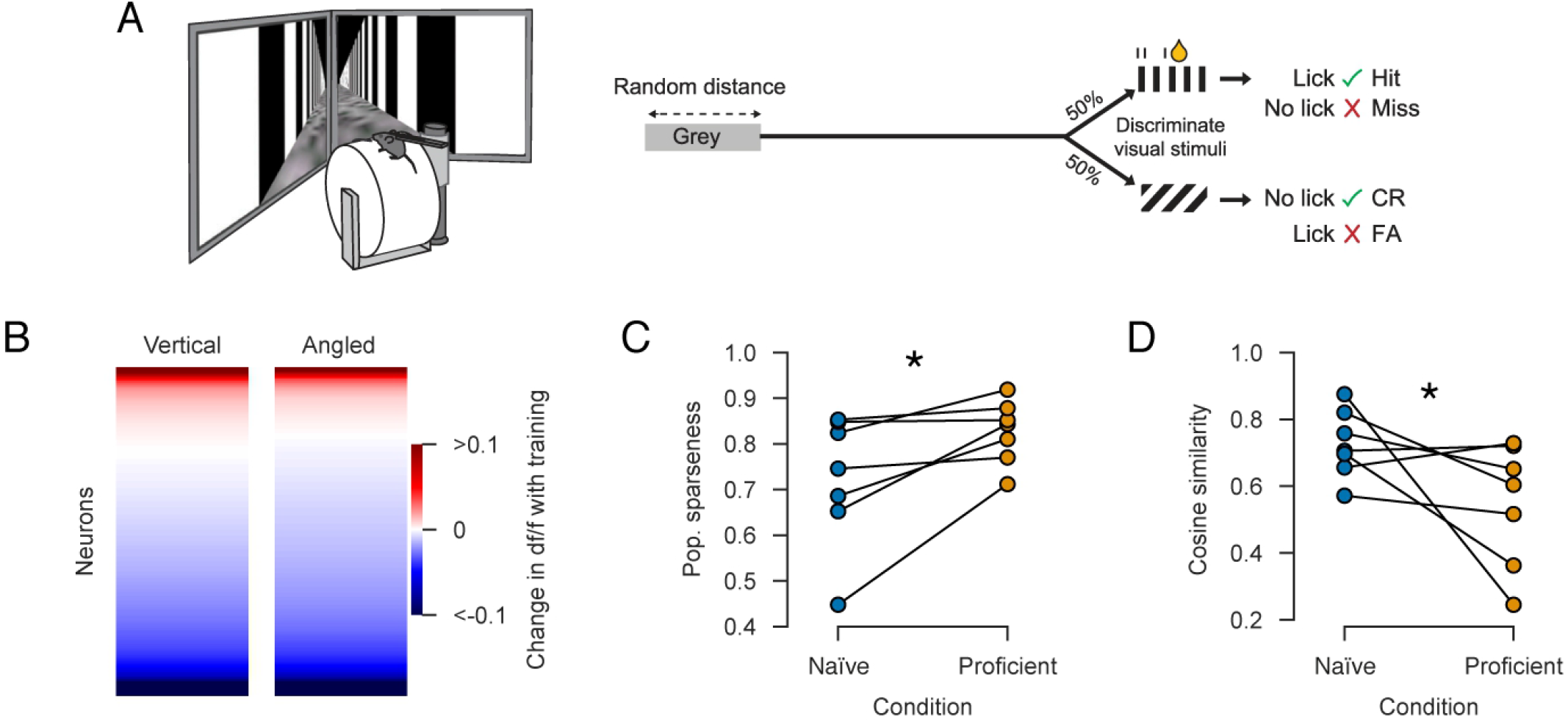
Analysis of data from a virtual reality discrimination task (Poort, et al., 2022). **A**, Mice ran on a wheel whose rotation was coupled to movement in a virtual corridor. Mice were trained to traverse the corridor and discriminate between a vertical and an angled grating. Licking the reward spout when a vertical grating was presented resulted in a water reward (figure adapted from Poort, et al., 2022). **B**, Change in the df/f response of neurons to the vertical (left) and angled (right) stimuli following training, pseudocolored so blue represents a decrease in activity after training, and red an increase. The response was calculated as the average df/f in a 0 – 1 s window relative to stimulus onset. Responses to both the vertical and angled gratings were suppressed following task training (p < 0.01; hierarchical linear mixed effects model; n = 1469 cells, N = 9 mice). **C**, Population sparseness increased with task training (p < 0.05, paired samples *t*-test, N = 7 mice). To ensure the accuracy of population level analyses, only mice with at least 100 recorded cells were included. **D**, Orthogonalization of population responses to the task stimuli, as shown by a decrease in their cosine similarity (p < 0.05, paired samples t-test, N = 7 mice).

## Appendix 1

To more deeply investigate our result that task training did not improve representational fidelity, we focused on coding of the motor-associated 45° and 90° stimuli, which require opposite behavioral contingencies in the task. We started by analyzing the coding properties of all recorded neurons individually. The mean response of a typical cell was lower after training, even when considering each neuron’s preferred motor-associated stimulus (Fig. A1A; hierarchical linear mixed effects model; p < 0.0001). Neuronal variability, assessed by the coefficient of variation of the response to each cell’s preferred stimulus, typically increased after training (Fig. A1B; hierarchical linear mixed effects, p < 0.0001) indicating that the decrease in mean response was not compensated by an equivalent decrease in standard deviation. Selectivity of neurons between the two motor-associated stimuli, assessed by a response index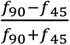, however, typically grew stronger, reflecting an increase in the percentage of neurons responding almost exclusively to one stimulus (Fig. A1C; hierarchical linear mixed effects model on the absolute value of the response index, p = 0.01). Finally, the d’ statistic, which measures how well a single neuron can distinguish between the two stimuli in the face of trial-to-trial variability, did not differ significantly between naïve and trained mice (Fig. A1D, note the log x-axis; hierarchical linear mixed effects model, p = 0.1), with small fraction of cells of very high d’ values (∼10) present in both cases. Thus, the effect of training on the average neuron was mixed: an increase in the difference between the task stimuli and an increase in coefficient of variation, leading to no systematic change in d’.

**Figure A1.**
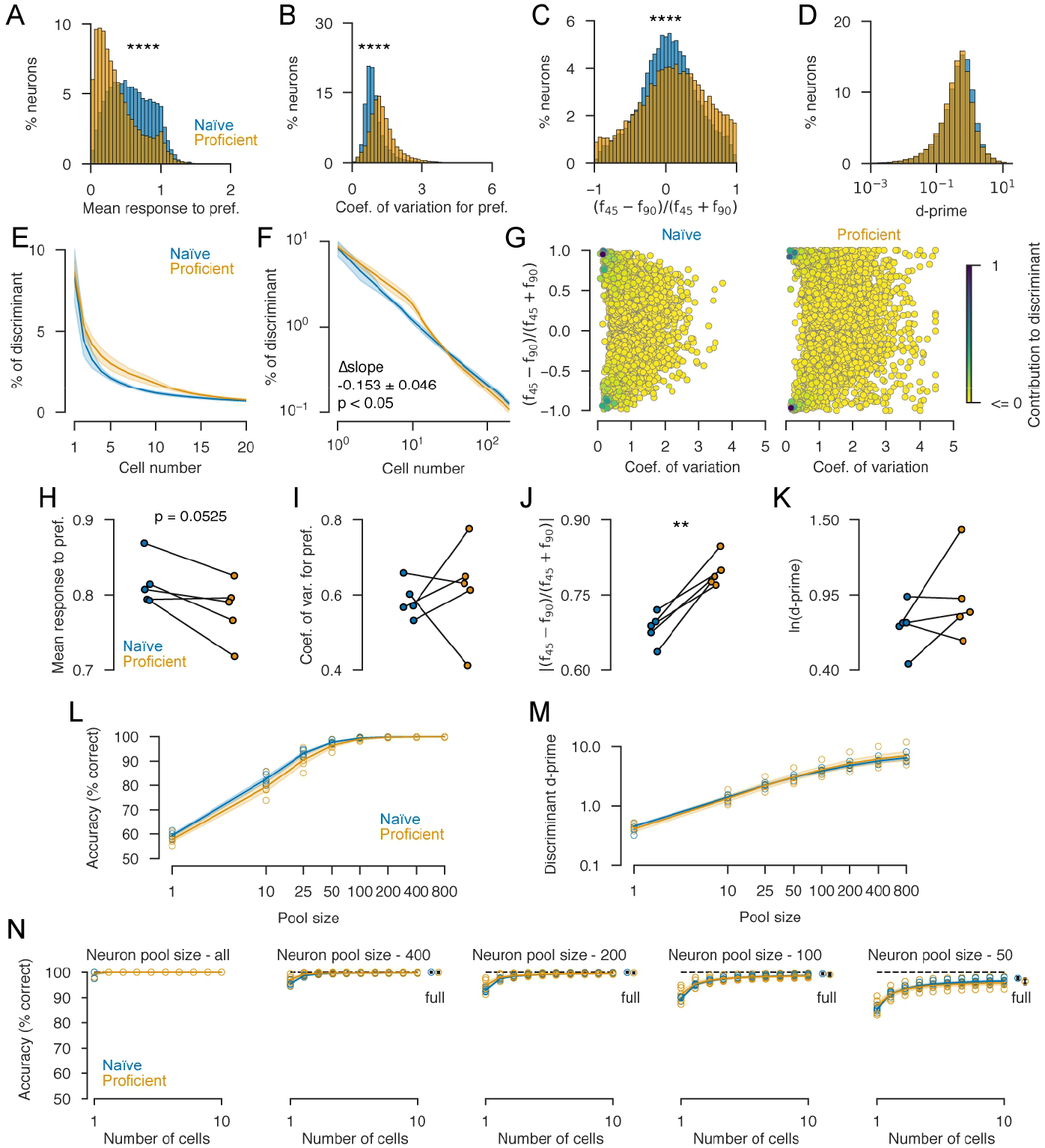
Deeper analysis of coding fidelity for motor-associated stimuli in naïve and trained mice. **A**, Histogram of mean response of each neuron to whichever of the two motor associated orientations (45° and 90°) drove it most strongly. Significance star: hierarchical linear mixed effects model. **B**, Histogram of coefficients of variation (standard deviation divided by mean) of each neuron’s responses to its preferred stimulus. **C**, Histogram of response index comparing activity evoked by the two stimuli, for all cells. **D**, Histogram of d’ discriminability for all cells (difference between means, divided by RMS standard deviation). For **A-D**, significance was assessed by a linear mixed effects model incorporating a random effect and slope for each mouse. **E**, Percentage of discriminant function accounted for by successive neurons, for an L2-regularized discriminant analysis classifier. Shading shows mean and SE over mice. **F**, same plot on a log-log scale. **G**, Analysis of cells contributing to discriminant function. Each circle represents a cell, in a position determined by its response index and coefficient of variation. Color represents percentage contribution to discriminant function. **H-K**, Average over neurons contributing to the decoder of the same statistics shown in (a-d), weighted by the neurons’ contributions to the discriminant function. **L**, Performance of a decoder trained on a randomly sub-selected pool of neurons, as a function of decoder size. No significant difference between naïve and proficient conditions was seen for any pool size. **M**, Similar plot measuring d’ of the discriminant function. Again, no difference was seen for any pool size. **N**, Accuracy of decoding from an optimal cell subset of neurons, selected from random pools of the indicated size. In no case was a significant difference between naïve and proficient conditions found. ****, p < 0.0001,

These changes did not affect the decodability of the stimuli, which was perfect for both naïve and proficient populations. To understand why, we analyzed the solution found by L2-regularized linear discriminant analysis, which computes a weighted sum of population activity (the “discriminant function”) with weights that maximize the reliable difference between the 45° and 90° stimuli. The decoder had 100% accuracy in all naïve and trained experiments when given access to the full ∼4000-cell population. To understand why changes in individual neuronal tuning did not affect performance, we investigated which neurons the decoder selected to base its decision on.

In both naïve and trained conditions, the decoder based its output on a sparse subset of neurons (Fig. A1E-G). To show this, we measured the percentage of the discriminant function accounted for by each neuron’s activity. The contribution of the recorded neurons to the discriminant function followed a power-law over the first ∼100 neurons (Fig. A1E, F): the proportion of the discriminant function accounted by the *n*^*th*^ neuron was approximately proportional to *n*^−*α*^, where the scaling exponent *α* was -0.760 ± 0.040 in naïve mice and -0.913 ± 0.071 in proficient mice, reflecting a small but significant increase in slope with training (p = 0.04, paired samples *t*-test). The single best neuron accounted for 8.4 ± 1.6% (naïve) or 8.6 ± 0.65% (proficient) of the discriminant function, and the top 20 neurons (∼0.5% of the recorded population) together accounted for 36.9 ± 3.1% and 46.7 ± 3.9% of the discriminant function (naïve and proficient; p < 0.05, paired samples *t*-test). The decoder thus based its decision on a highly sparse set of neurons, which became slightly but significantly sparser after training. Importantly, the L2-regularization approach that we used (unlike L1-based methods^72^) does not preferentially seek sparse weights; the fact that it nevertheless found them indicates that a sparse subset of neurons encoded the stimulus in a particularly advantageous manner.

The neurons selected by the decoder were strongly selective between the two task stimuli and had low variability (Fig. A1G), and in both naïve and proficient mice, there were enough such neurons to produce perfect decoding. The cells picked by the decoders again responded less in proficient than in naïve mice (p = 0.05, paired samples *t*-test) and showed higher selectivity (p = 0.002, paired samples *t*-test), but with no significant change in variability or d’ (p>0.05; Fig. A1H-K).

To further demonstrate how accurately this small set of neurons encoded the stimulus in both naïve and proficient conditions, we sequentially added neurons to our model based on their cross-validated performance (i.e., sequential feature selection), limiting the number of total neurons in our model to 10. Remarkably, decoding from just one optimally selected neuron yielded a cross-validated performance of 99.5 ± 0.5% in naïve mice, 99.6 ± 0.4% in proficient (Fig. A1N, left; p > 0.05, paired samples *t*-test).

Therefore, the 100% accuracy of stimulus decoding in naïve and trained mice arises because, in both conditions, a sparse subpopulation of cells encoded the stimulus extremely accurately. It remains possible, however, that a decoder denied access to these rare but exceptionally accurate neurons might work better in the trained condition. If so, this could constrain the decoding ability of downstream neurons in the brain that might have access to a limited subset of V1 axons, as well as explain the less-than-perfect decoding reported in previous experiments where a small population was recorded.

We therefore asked if a difference between naïve and trained decodability might appear for randomly selected cell pools, which will usually exclude the very best cells (Fig. A1L). When decoding from one randomly chosen neuron performance was 59.4 ± 0.6% in naïve mice, 57.8 ± 0.9% in proficient (p = 0.076, paired samples *t*-test), and increased in both cases to reach an asymptote of 100% at around 400 random neurons. For no pool size did we see a significant difference between naïve and proficient conditions. We also assessed decoder performance using the d’ of the discriminant function but again found no significant difference (Fig. A1M). We conclude that decoding fidelity does not increase following task training even for a decoder without access to the best neurons in the recorded population.

In a final attempt to find a decoder whose performance is better for proficient than naïve mice, we again picked an optimal sparse subset of each random cell pool in a sequential manner (Fig. A1N). In each case, decoding reaches asymptotic performance using just a few neurons, and again no significant difference was found between the naïve and trained conditions (p > 0.05 in all cases).

We conclude that while the structure of the V1 population code for orientation changes following task training, coding fidelity does not significantly improve in proficient mice, even after considering multiple methods to reveal such a difference.

## Appendix 2: mathematical results

### 2.1 Convex transformation of firing rates increases sparseness

Here we prove that applying a convex transformation to a neural population response vector increases its sparseness. Intuitively, the argument works as follows. Sparseness measures the degree to which a small number of neurons fire more than the mean firing rate. Applying a convex transformation causes a disproportionate boost in the firing rate of these few highly active neurons, increasing the sparseness of the population response.

Formally, we will prove that this holds for a wide family of sparseness metrics, which includes those described by Treves and Rolls and Willmore and Tolhurst^39,40^ as a special case corresponding to *k*(*x*) = *x*^2^.

#### Theorem

*Let k*(*x*) *be a convex function. Let* {*x*_*i*_: *i* = 1 … *N*} *be a finite set of non-negative real numbers. We define the sparseness measure*

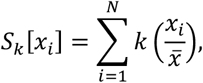

*Where*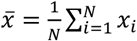. *Let g be a convex non-decreasing function with g*(0) = 0, *and write y* = *g*(*x*). *Then*

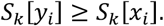

**Proof**. For any scalar *a, S*_*k*_[*x*_*i*_] = *S*_*k*_[*ax*_*i*_]. So, without loss of generality, we can rescale *x* and *g* so that 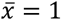 and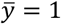. After this rescaling,

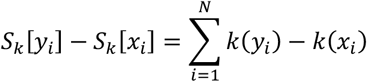

Now because ∑_*i*_ *x*_*i*_ = ∑_*i*_ *g*(*x*_*i*_), and *g* is continuous, there must exist an *x*_0_ with *g*(*x*_0_) = *x*_0_. Because *g* is convex and *g*(0) = 0, *x*_*i*_ ≥ *x*_0_ implies *y*_*i*_ ≥ *x*_*i*_, and *x*_*i*_ ≤ *x*_0_ implies *y*_*i*_ ≤ *x*_*i*_. Let *d* be a subgradient of *k* at *x*_0_, so if either *a* ≥ *b* ≥ *x*_0_ or *a* ≤ *b* ≤ *x*_0_, then *k*(*a*) − *k*(*b*) ≥ *d*(*a* − *b*). If *x*_*i*_ ≥ *x*_0_ then *y*_*i*_ ≥ *x*_*i*_ ≥ *x*_0_ and if *x*_*i*_ ≤ *x*_0_ then *y*_*i*_ ≤ *x*_*i*_ ≤ *x*_0_. For all *i* one of these two conditions is true so *k*(*y*_*i*_) − *k*(*x*_*i*_) ≥ *d*(*y*_*i*_ − *x*_*i*_). Thus 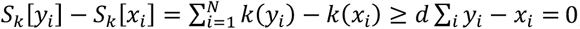, as we have rescaled so that ∑_*i*_ *x*_*i*_ = ∑_*i*_ *y*_*i*_. Thus, *S*_*k*_[*y*_*i*_] ≥ *S*_*k*_[*x*_*i*_] and the theorem is proved.

### 2.2 Connection between sparsity and orthogonality

The fact that sparse codes are more orthogonal was noted many years ago ^42^. If neuronal activity is approximated as binary, the connection between sparseness and orthogonality can be easily understood. In a sparse code most do not respond to most stimuli, so the number of neurons responding to both of two stimuli will be very small. Because the cosine angle of two binary response vectors depends on the number of neurons driven by both stimuli, it will be smallest for sparse codes.

We now demonstrate this idea for continuous-rate codes. Let the response of cell *i* to a first stimulus be *x*_*i*_ and its response to the second be *y*_*i*_, where *i* = 1 … *N* indexes the cell number. We assume that the response of a cell to one stimulus is independent of its response to the other. Thus, we can consider *x*_*i*_ and *y*_*i*_ to be independently drawn from a probability distribution with mean 𝔼 [*x*_*i*_] = *μ* and second moment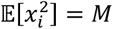. By independence of the two rate vec-tors,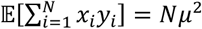. In the limit of large population size, the expected cosine angle between the two vectors con-verges to

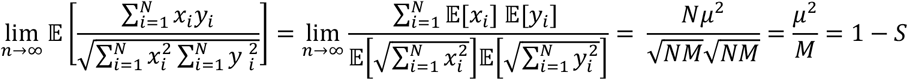

Where *S* is the Treves-Rolls sparseness measure. Thus, we see orthogonality is negatively related to sparseness, with maximally sparse codes having a cosine angle close to 0, and minimally sparse codes having a cosine angle close to 1. As an example, consider a binary code in which 1 out of K cells are activated by either stimulus, firing with rate f. Then 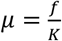 and 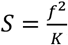,so the expected cosine angle is 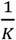, indicating that sparser codes are more orthogonal.

### 2.3 Subtractive inhibition can mediate convex rate transformation

To show increased activity of inhibitory neurons could cause a convex transformation of firing rates, we consider the effect of subtractive inhibition on a firing-rate model neuron with a sigmoid activation function. We model the cell’s firing rate as σ (*x* − Δ), where *x* is the excitatory input current, Δ is the inhibitory input current, and 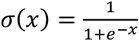 is the logistic sigmoid.

To see how subtractive inhibition transforms firing rates, we need to find a function *g* _Δ_ (*f*) such that *g*_Δ_ (*σ* (*x*)) = *σ* (*x* − Δ). This function is *g*_Δ_ (*f*) = *σ* (*σ*^−1^(*f*) − Δ), where 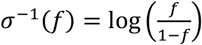 is the inverse function of the logistic sig-moid. We thus have

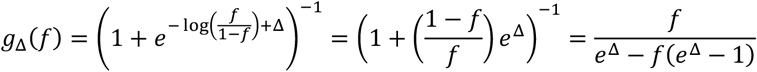

This function is convex over the range of possible firing rates *f* = 0 … 1, as can be seen from the fact that its second derivative is positive in this range. Its convexity increases with greater inhibition strength, as shown in the plot below.

**Figure A2.**
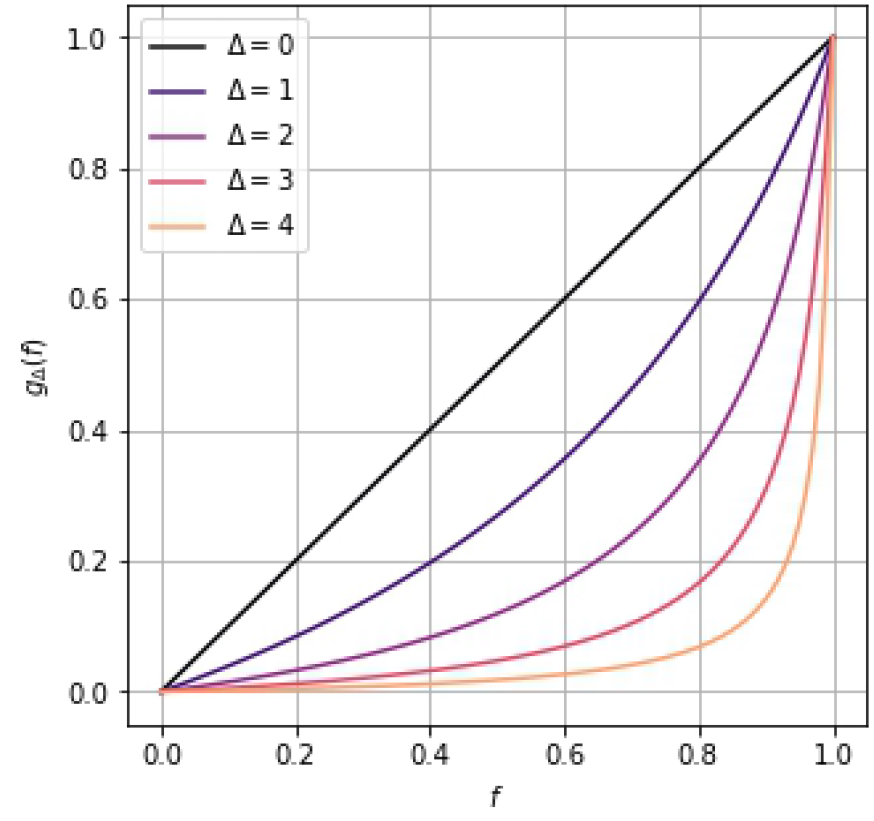
Convex transformation of firing rates by subtractive inhibition, for different levels of inhibition strength Δ.

